# Monoclonal nanobodies alter the activity and assembly of the yeast vacuolar H^+^-ATPase

**DOI:** 10.1101/2025.01.10.632502

**Authors:** Kassidy Knight, Jun Bae Park, Rebecca A. Oot, Md. Murad Khan, Soung-Hun Roh, Stephan Wilkens

## Abstract

The vacuolar ATPase (V-ATPase; V_1_V_o_) is a multi-subunit rotary nanomotor proton pump that acidifies organelles in virtually all eukaryotic cells, and extracellular spaces in some specialized tissues of higher organisms. Evidence suggests that metastatic breast cancers mislocalize V-ATPase to the plasma membrane to promote cell survival and facilitate metastasis, making the V-ATPase a potential drug target. We have generated a library of camelid single-domain antibodies (Nanobodies; Nbs) against lipid-nanodisc reconstituted yeast V-ATPase V_o_ proton channel subcomplex. Here, we present an in-depth characterization of three anti-V_o_ Nbs using biochemical and biophysical *in vitro* experiments. We find that the Nbs bind V_o_ with high affinity, with one Nb inhibiting holoenzyme activity and another one preventing enzyme assembly. Using cryoEM, we find that two of the Nbs bind the *c* subunit ring of the V_o_ on the lumen side of the complex. Additionally, we show that one of the Nbs raised against yeast V_o_ can pull down human V-ATPase (*Hs*V_1_V_o_). Our research demonstrates Nb versatility to target and modulate the activity of the V-ATPase, and highlights the potential for future therapeutic Nb development.

## Introduction

The vacuolar ATPase (V-ATPase; V_1_V_o_-ATPase) is a multi-subunit proton pump present in virtually all eukaryotes that works to acidify the lumen of organelles (Eaton et al, 2021; Forgac, 2007; Futai et al, 2000; Kane, 2007; Marshansky, 2022). In special cases, the V-ATPase is also responsible for the acidification of extracellular space in specific cell types such as in osteoclasts and renal intercalated cells (Sun-Wada et al, 2003; Toei et al, 2010). V-ATPases play key role in organelle pH and ion homeostasis, which is required for protein trafficking/degradation in the exocytic and endocytic pathways (Stasic et al, 2019). V-ATPase function is also important in autophagy (Han et al, 2022; Xu et al, 2022), phagocytosis (Hooper et al, 2022), mTOR and Notch signaling (Xu et al, 2012; Yan et al, 2009; Zoncu et al, 2011), inflammation (Rao et al, 2019), hormone secretion (Sun-Wada, 2006), and neurotransmitter release (Vavassori & Mayer, 2014).

The V-ATPase consists of two subcomplexes: a cytosolic V_1_-ATPase and a membrane bound V_o_ proton channel. In yeast, a well characterized model system for the V-ATPase of higher organisms, including human, the two subcomplexes contain 15 different polypeptides, with V_1_ and V_o_ made of A_3_B_3_(C)DE_3_FG_3_H and *ac_8_c’c”def*Voa1p, respectively (**Fig. 1**). The proteolipid isoform *c’* is only found in some fungi, including yeast. The V_o_ of higher organisms contains two additional polypeptides, Ac45 (ATP6AP1) and the C-terminal transmembrane domain (TMD) of the prorenin receptor (ATP6AP2), with the C-terminal TMD of Ac45 being related to the C-terminal TMD of the yeast V_o_ assembly factor, Voa1p (Jansen et al, 2016).

**Figure 1:**
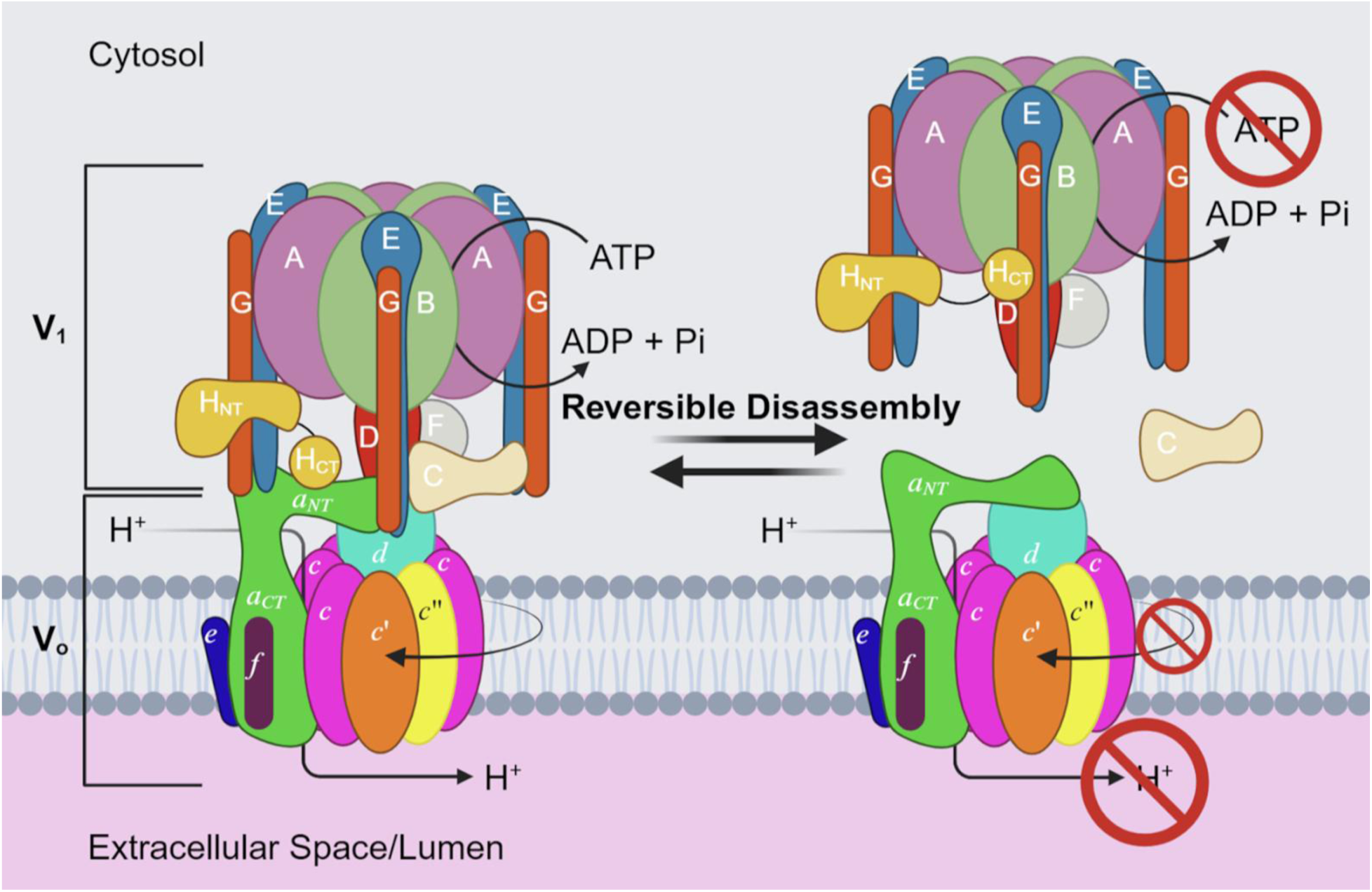
Structure and reversible disassembly of the yeast V-ATPase. The V-ATPase is composed of two subcomplexes: A cytosolic V_1_ (A_3_B_3_CDE_3_FG_3_H) and membrane integral V_o_ (*ac*_8_*c*’*c*”*def*). The two subcomplexes can be found assembled in holo V-ATPase (left), and disengaged from each other (right). When assembled, ATP hydrolysis on V_1_ is paired to proton translocation through V_o_. During regulation by reversible disassembly, when V_1_ is released from V_o_, the C subunit dissociates into the cytosol and both subcomplexes become autoinhibited.

Proton pumping by the V-ATPase is achieved by a rotary mechanism: ATP hydrolysis in three catalytic sites on V_1_ drives rotation of the *central rotor* made of subunits D,F,*d*, and the ring of *c,c’,c”* subunits (*c*-ring) relative to the *stator* complex consisting of V_o_ subunits *aef* and the rest of the V_1_ (A_3_B_3_E_3_G_3_HC). Upon rotation of the *c*-ring, protons are transferred from the cytosol into the lumen of the subcellular compartment (or the extracellular space in case of V-ATPases in the plasma membrane of specialized animal cells) through two half-channels located within subunit *a*’s C-terminal domain (*a*_CT_) (Mazhab-Jafari et al, 2016; Roh et al, 2020; Roh et al, 2018). Three peripheral stalks made of heterodimers of subunits E and G connect the A_3_B_3_ catalytic hexamer of the V_1_ to the single-copy subunits H and C and the N-terminal domain of subunit *a* (*a*_NT_) to withstand the torque of rotary catalysis. Cryo electron microscopy (cryoEM) structures of V-ATPases from yeast (Khan et al, 2022; Vasanthakumar et al, 2022; Zhao et al, 2015) and mammals (Abbas et al, 2020; Wang et al, 2021; Wang et al, 2020) have captured the enzyme in three *rotary states* referred to as states 1-3, with the different states showing the central DF*dc*-ring rotor in three rotary positions separated by 120°.

V-ATPase’s activity is tightly regulated, most prominently by a mechanism referred to as *reversible disassembly*. When V-ATPase activity is not required, the enzyme reversibly dissociates into V_1_-ATPase and V_o_ proton channel subcomplexes (Kane, 1995; Sumner et al, 1995), with both subcomplexes becoming autoinhibited and locked in rotary states 2 and 3, respectively, to prevent unregulated re-assembly (Couoh-Cardel et al, 2015; Merzendorfer et al, 1997; Oot, 2016; Oot et al, 2017; Parra et al, 2000; Sharma et al, 2018; Vasanthakumar et al, 2022; Zhang et al, 1994) (**Fig 1**). The process, while best characterized in yeast, is conserved in mammalian cells, though the signals that govern the assembly state of the V-ATPase vary.

Whereas enzyme assembly on yeast vacuoles is driven by the presence of glucose (Parra & Kane, 1998), assembly on mammalian lysosomes is increased under conditions of starvation to replenish nutrients (Ratto et al, 2022). In cultured neurons, synaptic vesicle-associated V-ATPases disassemble once neurotransmitter loading is complete and reassemble after SV fusion with the presynaptic membrane, with the luminal or extracellular pH being one of the signals driving assembly/disassembly (Bodzeta et al, 2017). Studies in yeast have shown that efficient V-ATPase assembly requires the heterotrimeric RAVE complex (Seol et al, 2001; Smardon et al, 2014), while disassembly is facilitated by the TLDc domain protein Oxr1p (Khan et al, 2022; Khan & Wilkens, 2024).

Several subunits of the V-ATPase are expressed as multiple isoforms. In yeast, V_o_ subunit *a* is expressed as two isoforms, Vph1p and Stv1p, forming V-ATPase complexes that localize to the vacuole and Golgi, respectively (Manolson et al, 1994). In mammals, subunit *a* has four isoforms (*a*1-4) that have been found enriched in different tissues or compartments, with *a*1 on lysosomes and synaptic vesicles, *a*2 on the Golgi and early endosomes, *a*3 on secretory lysosomes and the ruffled membrane of bone osteoclasts and *a*4 on the apical membrane of ⍺ intercalated cells in the kidney (Toei et al, 2010). As diseases linked to V-ATPase are frequently associated with enzyme populations containing specific subunit isoform combinations that are located on specialized membranes in the cell (Frattini et al, 2000; Jansen et al, 2016; Karet, 1999; Kornak et al, 2008; Smith et al, 2000), this offers opportunities for therapeutic targeting of disease causing or promoting complexes. For example, isoform *a*3 and *a*4 containing V-ATPases are frequently found mislocalized to the plasma membrane of highly invasive cancer cells, presumably to extrude metabolic acid, which in turn is thought to help activate metalloproteases that facilitate cell migration (Collins & Forgac, 2020; Cotter et al, 2016; Stransky et al, 2016).

Here we characterize a set of single-domain camelid antibodies (Nanobodies; Nbs) raised against lipid nanodisc-reconstituted yeast V_o_ subcomplex (V_o_ND). We show that some Nbs inhibit the activity of holo V-ATPase while another prevents enzyme assembly from V_1_ and V_o_ subcomplexes. Structures obtained by cryoEM show that Nb-mediated inhibition of the holoenzyme is due to binding of Nbs on the luminal side of the *c*-ring, with the Nbs likely hindering *c*-ring rotation past *a*_CT_. On the other hand, inhibition of assembly is caused by Nb binding to the V_o_ *d* subunit at a location that appears to interfere with V_1_ binding due to steric clashing with the central rotor. Moreover, we found that one of the Nbs that binds the luminal side of the *c*-ring was able to pull down purified human V-ATPase, demonstrating the potential of ⍺-yeast V-ATPase Nbs in targeting human V-ATPase for diagnostic or therapeutic purposes.

## Results

### Nanobody Expression and Purification

To explore new means of selectively modulating V-ATPase activity either depending on the enzyme’s location in the cell, or via influencing complex’s assembly state, we generated a library of single-domain camelid antibodies, or Nanobodies, directed against the purified and lipid nanodisc reconstituted yeast V_o_ subcomplex (V_o_ND). Yeast V_o_ subcomplex was affinity purified via a calmodulin binding peptide fused to the C-terminus of the Vph1p isoform of subunit *a* and reconstituted into lipid nanodiscs as described previously (Couoh-Cardel et al, 2015; Stam & Wilkens, 2016) (**Fig. 2A**). Immunization, cDNA library construction, antigen specific Nb selection via phage display and ELISA were carried out by the VIB Nanobody facility at the Vrije Universiteit Brussel, Brussels, Belgium. Specificity was determined by screening against V_o_ND, with empty nanodiscs and membrane scaffold protein (MSP) as negative controls. Following Llama immunization and Nb selection, we received 94 Nb clones that were distributed over 38 groups based on the sequence similarity of the primary antigen binding loop (complementary determining region 3; CDR3). Of the 94 Nbs, 35 contained internal Amber or Opal stop codons, and these Nbs were excluded from the initial analysis. We then selected Nbs from different CDR3 groups, based on the highest signal from the ELISA screen provided by the Nanobody facility, for further characterization. Nbs were tested for their impact on V-ATPase’s ATP hydrolysis activity and *in vitro* enzyme assembly from purified components (**Fig 2A**). Nb clones were expressed in *Escherichia coli*, with a yield of 8-13 mg purified Nb per liter of culture (**Fig. 2B; Appendix Fig. S1**).

**Figure 2:**
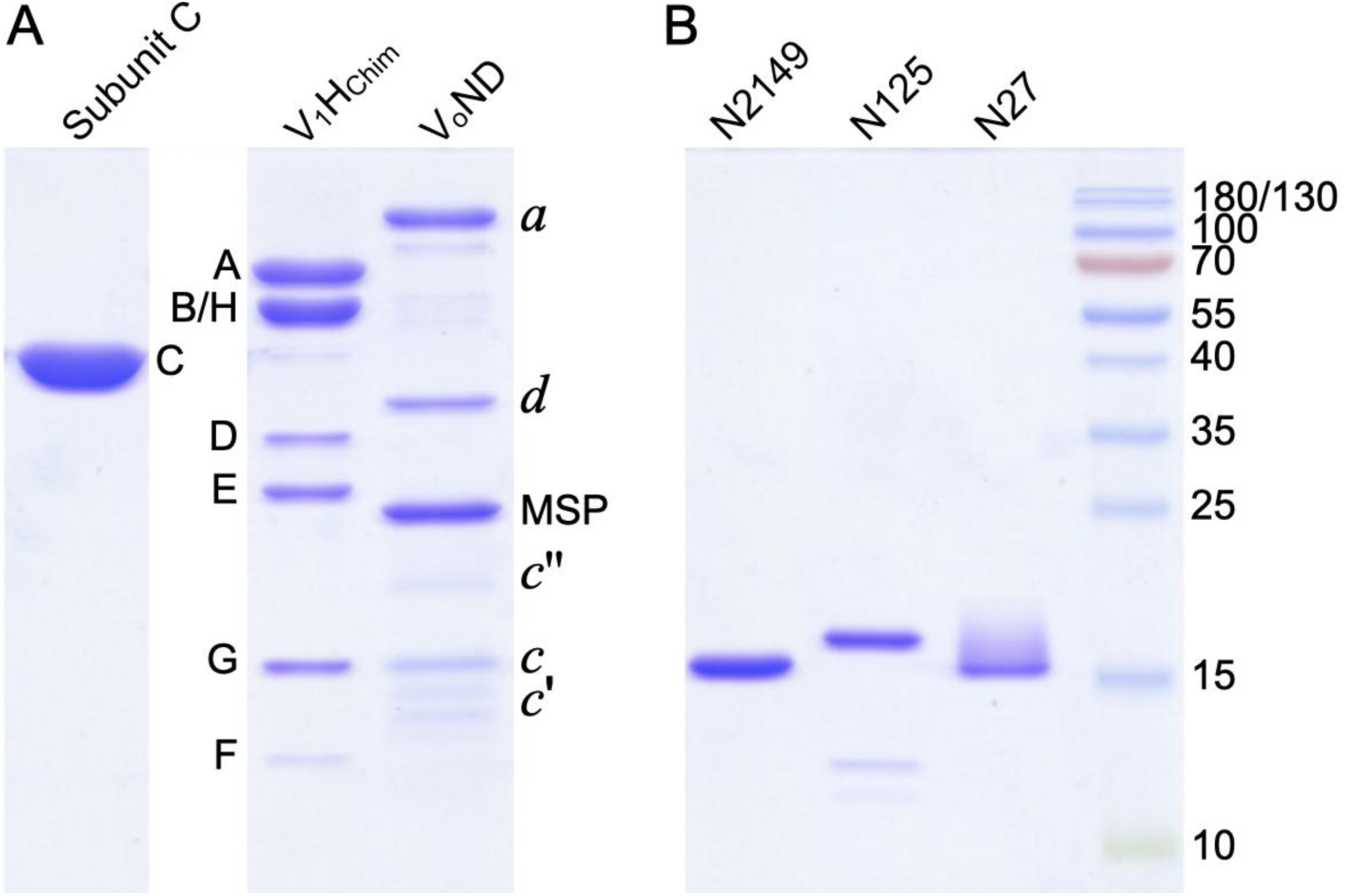
SDS-PAGE of purified yeast V-ATPase components and Nbs. From left to right: Recombinant yeast subunit C (1.2 µg), V_1_(H_chim_) (7.4 µg), V_o_ND (3.3 µg), Nbs N2149 (2.3 µg), N125 (1.5 µg), N27 (3 µg). V_1_ΔH (used to prepare V_1_(H_chim_)) and V_o_ND were purified from yeast, and C, H_chim_ and Nbs were expressed in *E. coli*.

### *In vitro* assembly of yeast V-ATPase and Nanobody characterization

Despite a negative ΔG for the overall process, purified wild type V_1_ and V_o_ (plus recombinant subunit C) do not assemble spontaneously *in vitro*, presumably due to the large activation barrier for overcoming autoinhibited conformations of the two subcomplexes (Oot et al, 2017; Wilkens et al, 2023). We previously showed that replacing the inhibitory V_1_-H subunit with a chimeric polypeptide containing yeast N-terminal and human C-terminal domains (H_chim_) relieves V_1_ autoinhibition (Oot, 2016; Sharma et al, 2019). The V_1_ complex containing H_chim_ (referred to as V_1_(H_chim_); **Fig. 2A**) is generated by adding recombinant H_chim_ to V_1_ purified from a yeast strain deleted for subunit H (V_1_ΔH). We further showed that the V_1_(H_chim_) together with recombinant subunit C (**Fig. 2A**) readily binds V_o_ND to form holo V-ATPase (hereafter referred to as V_1_(H_chim_)V_o_ND) (Khan et al, 2022; Sharma et al, 2019). The *in vitro* assembled V_1_H_chim_V_o_ND has robust magnesium dependent ATPase (MgATPase) activity, which is sensitive to the V-ATPase specific inhibitor, Concanamycin A (ConA), indicating that V_1_ and V_o_ are functionally coupled (Khan et al, 2022; Sharma et al, 2019). To test the impact of the anti-V_o_ND Nbs on V-ATPase activity and enzyme assembly, we either incubated purified Nb with *in vitro* assembled holo V-ATPase followed by ATPase activity measurement (*activity assay*), or we pre-incubated purified V_o_ND with Nb before assembly and then performed ATPase activity assays (*assembly assay*). Based on preliminary experiments with a set of nine Nbs from eight CDR3 groups, we chose three Nbs from different CDR3 groups for a more detailed analysis: 2WVA7, 1WVA25, and 2WVA149, in the following referred to as N27, N125, and N2149, respectively (**Fig 2B**).

Measurements are at least two technical replicates each from three independent purifications of V-ATPase and two independent purifications of Nbs. Error bars are SEM. P values were calculated using an unpaired t test assuming different SD, with ns>0.05, *≤0.05, ***≤0.001, and ****≤0.0001. The specific V-ATPase inhibitor ConcanamycinA (ConA) was used as a control.

### Nanobody mediated impact on V-ATPase activity

To determine the Nbs’ impact on the ATPase activity of the assembled enzyme, each Nb was added individually to *in vitro* assembled V_1_(H_chim_)V_o_ND and incubated for a minimum of 30 minutes at room temperature. The specific ATPase activity of Nb bound V_1_(H_chim_)V_o_ND was then measured using an ATP regenerating assay, with V_1_(H_chim_)V_o_ND (no added Nb) as a control.

The average specific ATP hydrolysis activity of *in vitro* reconstituted V_1_(H_chim_)V_o_ND was 6.7 ± 0.5µmol/(min×mg). Upon addition of 200 nM of the specific V-ATPase inhibitor ConA, the activity was reduced to 0.6 ± 0.03 µmol/(min×mg) (**Fig. 3A**). We found that N27 needs to be added in excess over V_o_ND to achieve full inhibition and we therefore used a 24-fold molar excess for activity measurements. Under these conditions, addition of N27 resulted in close to complete loss of ATPase activity (84%), with the activity further reduced by the addition of ConA. On the other hand, incubating the enzyme with N125 (4.5 molar excess) or N2149 (7.6 molar excess) had no significant impact on the activity (**Fig. 3A**). Overall, we found that whereas N27 inhibited the ATPase activity of intact V_1_(H_chim_)V_o_ND almost completely, N125 and N2149 had no significant effect on the activity of the *in vitro* assembled complex.

**Figure 3:**
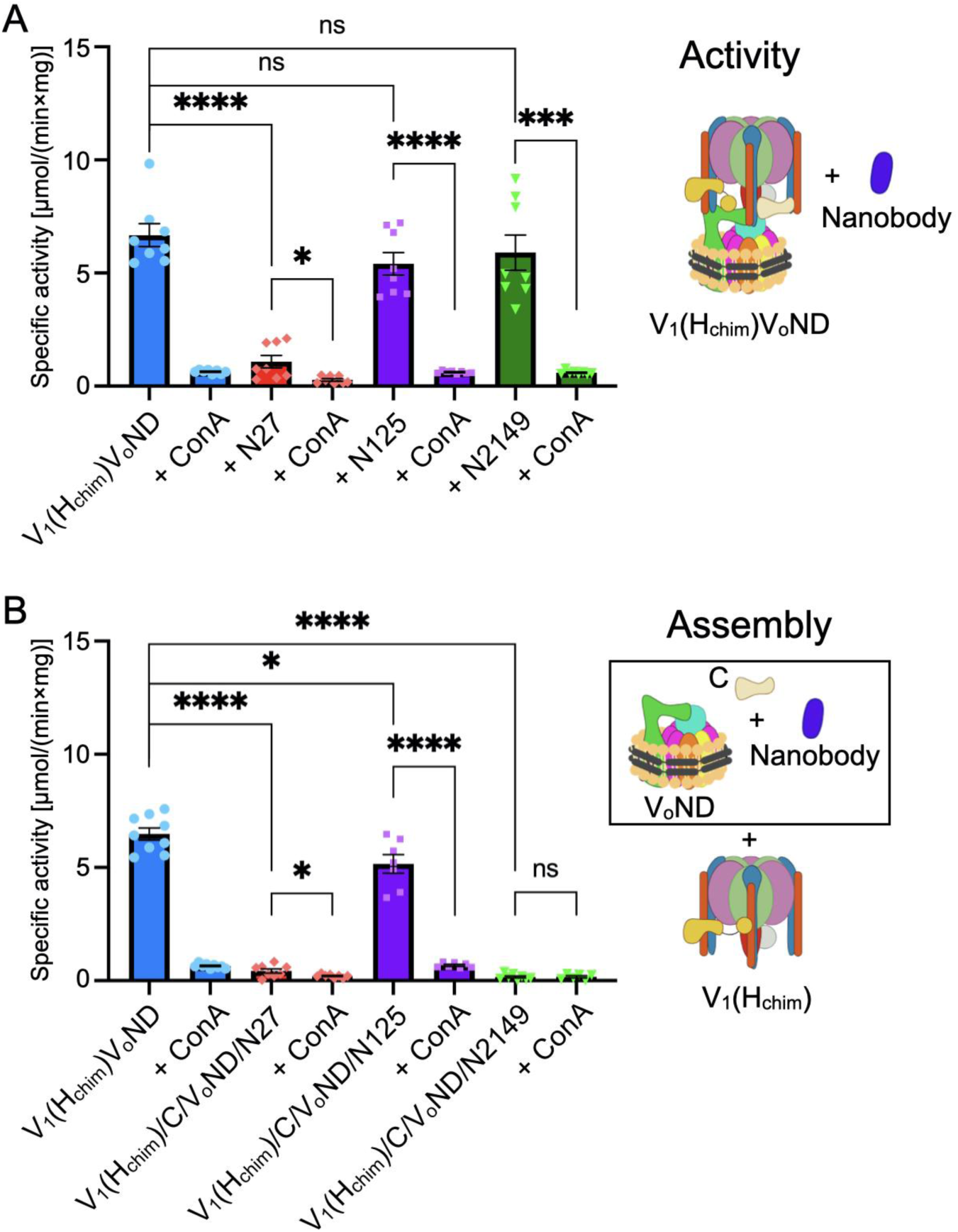
Activity assay and Assembly assay. **A** V_1_(H_chim_)V_o_ND (average specific activity of 6.8 µmol/(min×mg)) was incubated with purified Nbs N27, N125 and N2149 before measuring Mg^2+^ dependent ATPase activity in an ATP regenerating assay at 37 °C. **B** V_o_ND was incubated with purified Nbs N27, N125 and N2149 and incubated for 30 min at room temperature. Nb-bound V_o_ND was then incubated with a 1:1 molar ratio of V_1_H_chim_ and a 3- fold molar excess of subunit C for two hours at room temperature or overnight at 18 °C.

### Nanobody mediated impact on V-ATPase assembly

As the Nbs were raised against the V_o_ND subcomplex, we next wanted to establish whether the Nbs could interfere with V_1_(H_chim_)V_o_ND assembly. To do so, Nbs were incubated with purified V_o_ND prior to *in vitro* assembly. Nb-bound V_o_ND, V_1_H_chim_ and subunit C were incubated at room temperature for 2 h or 18 °C overnight before measurement of ATPase activity, with V_1_(H_chim_)V_o_ND assembled in absence of Nb as a control. Of note, holo V-ATPase activity is characterized by a linear slope over time in an ATP regenerating assay (**Appendix Fig. S2A**). On the other hand, the ATPase activity of V_1_H_chim_, which is used to assemble the holoenzyme, has robust ATPase activity, but the activity quickly decays due to MgADP inhibition (**Appendix Fig S2B**). Thus, if the addition of a Nb were to prevent assembly, the expectation is that the ATPase activity of the assembly mix will resemble the activity of V_1_H_chim_. If, however, assembly is successful, but the resulting holoenzyme is inactive due to Nb binding to V_o_ND, the resulting activity profile should be linear, with the slope depending on the inhibitory potency of the Nb. Unsurprisingly, when pre-incubating V_o_ND with N27 before adding V_1_H_chim_, very little activity is obtained (**Fig. 3B**), consistent with N27’s ability to inhibit the holo V-ATPase (**Fig. 3A**). When inspecting the activity trace of the assembly mixture containing N27, however, we did not observe any activity due to unassembled V_1_H_chim_ (**Appendix Fig. S2C**), suggesting that binding of N27 to V_o_ND did not prevent assembly of the holoenzyme. This interpretation is consistent with earlier experiments in yeast that showed that V-ATPase assembly in the vacuole occurs even in presence of the specific V-ATPase inhibitor, ConA, which is known to bind V_o_ (Parra & Kane, 1998). N125 reduced the activity by ∼20% compared to the control, with the remaining ∼80% of the activity demonstrating ConA sensitivity, indicating that N125 did not inhibit assembly of V_1_H_chim_ and V_o_ND under the experimental conditions **(Fig. 3B)**. Whereas N2149 had no significant effect on the activity of the assembled enzyme (**Fig. 3A**), the activity profile of the N2149 containing assembly mix indicated the presence of an early burst of activity (**Appendix Fig. S2D**), consistent with the presence of unassembled V_1_H_chim_. The slope past the early activity burst, however, resembled the residual activity of V_1_H_chim_ alone, indicating that binding of N2149 to V_o_ND prevented holoenzyme assembly **(Fig. 3B)**. Furthermore, while negative stain TEM of the control assembly mix containing V_o_ND, V_1_H_chim_ and C showed a significant proportion of assembled V-ATPase complexes (**Appendix Fig. S3A**), images of the assembly mix containing N2149 showed only unassembled V_1_H_chim_ and V_o_ND particles (**Appendix Fig. S3B**), consistent with the lack of ConA sensitive holoenzyme activity (**Fig. 3B**). Overall, we found that despite being inhibitory, N27 did not seem to prevent assembly of the holoenzyme, whereas N2149 binding to V_o_ND, which has no effect on activity of the holoenzyme complex, prevented assembly with V_1_H_chim_ and subunit C. N125 on the other hand had little effect on the ability of V_o_ND to form an active complex.

### Nanobodies bind V_o_ND with high affinity

To determine the affinity of the Nbs for V_o_ND, we quantified the binding kinetics using biolayer interferometry (BLI). Here, each nanobody was immobilized on Ni-NTA sensors via a C-terminal 6×His tag, and the kinetics of V_o_ND association and dissociation was monitored over time. The experiment was carried out with a series of V_o_ND concentrations ranging from 0.083 nM to 60 nM for each Nb, and non-specific binding to empty sensors and nanobody-only dissociation controls were subtracted using a 16-biosensor setup. Global fitting of the association and dissociation traces revealed that all three Nbs bind lipid nanodisc reconstituted V_o_ with pM affinities (**Fig. 4**). As there was essentially no significant dissociation of V_o_ND from immobilized N27 over the observation time period of 1 h, the K_d_ for N27 was given as <1 pM, which is the lower limit of the algorithm used for data fitting **(Fig 4A,D)**. The average binding affinities for N125 and N2149 were determined to be 13 ± 1 and 90 ± 23 pM, respectively **(Fig 4B,C,D)**.

**Figure 4:**
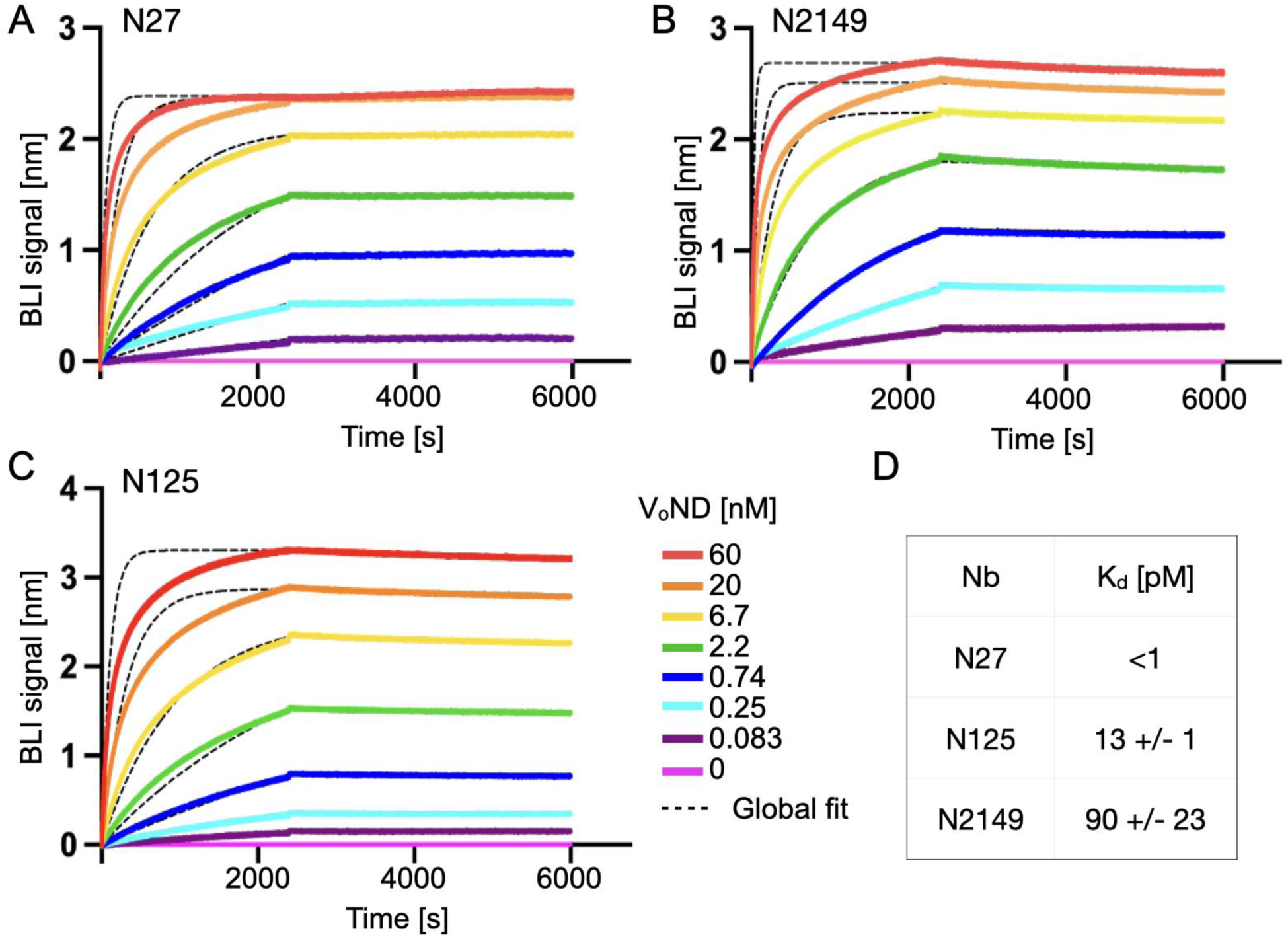
Affinity between V_o_ND and Nbs as determined by BLI. **A-C** C-terminally His-tagged N27, N125, and N2149 were immobilized on NiNTA sensors and V_o_ND binding was measured over a concentration range of 0.083 to 60 nM. **D** K_d_s were obtained by global fitting of the association and dissociation traces. Fits are represented by the dashed lines.

### Nb epitope mapping using cryo-EM structure determination

Because the Nbs were raised and screened against the multi-subunit yeast V_o_ND subcomplex, we wished to determine which subunits N27, N125 and N2149 bind to. To determine the Nbs’ epitopes, we used cryoEM structure determination.

### Epitope of N27

From the observations that N27 binding to V_o_ND occurred with pM affinity and that full inhibition of enzyme activity required a large molar excess of N27, we speculated that N27 may bind the *c* subunit as this is the only subunit in V_o_ that is present in multiple (eight) copies. We therefore incubated *in vitro* assembled V_1_(H_chim_)V_o_ND with a 16-fold molar excess of N27 before the mixture was vitrified for cryoEM imaging. A dataset of 3215 movies recorded on a 200 kV cryoEM instrument was analyzed using cryoSPARC (Punjani et al, 2017) and RELION (Kimanius et al, 2021) software packages.

2-D classification and *ab initio* 3-D reconstructions of a dataset of ∼500,000 particles using cryoSPARC revealed presence of holo V_1_(H_chim_)V_o_ND complexes (∼30%) as well as V_o_ND (∼8%) and V_1_ (∼30%) subcomplexes (**Appendix Fig. S4**). The map of the V_1_ subcomplex, which was obtained at a resolution of ∼3.2 Å, resembles the previously described structure of V_1_ bound to Oxr1p and subunit C (V_1_(C)Oxr1p) (Khan et al, 2022). The V_1_(C)Oxr1p subcomplex co-purifies from yeast cells with the V_1_(ΔH) complex used for preparing V_1_H_chim_, which is required for *in vitro* reconstitution of the holoenzyme (Khan et al, 2022; Sharma et al, 2019). For *in vitro* reconstitution, V_1_H_chim_ (containing varying amounts of V_1_(C)Oxr1p) and V_o_ND were mixed at a 1:1 molar ratio, but since V_1_(C)Oxr1p does not assemble with V_o_ND (Khan et al, 2022; Khan & Wilkens, 2024), this may explain the presence of free V_o_ND bound to N27 in the assembly mix analyzed by cryoEM. The 3-D reconstruction of N27 bound V_1_(H_chim_)V_o_ND as obtained using cryoSPARC indicated some heterogeneity in the population of N27-bound holoenzymes, and we therefore analyzed the dataset in more detail focusing on rotary states and substates of the N27 bound V_o_ND and V_1_(H_chim_)V_o_ND complexes. From this analysis, we obtained a ∼3.3 Å resolution reconstruction of N27 bound to V_o_ND in the canonical rotary state 3 (**Fig. 5**; **Appendix Fig. S5**), and reconstructions of the N27 bound holoenzyme in rotary state 1 (22%, 4.4 Å), the previously described rotary state 3’ (Roh et al, 2020) (45%, 3.8 Å; here referred to state 3^+18^) and a hitherto not observed rotary state here referred to as state 3^-18^ (33%, 4.3 Å) (**Fig. 6; Appendix Fig. S6**). The structure of the N27 bound V_o_ND subcomplex will be described in the following, with the structures of N27 bound holoenzymes further below.

**Figure 5:**
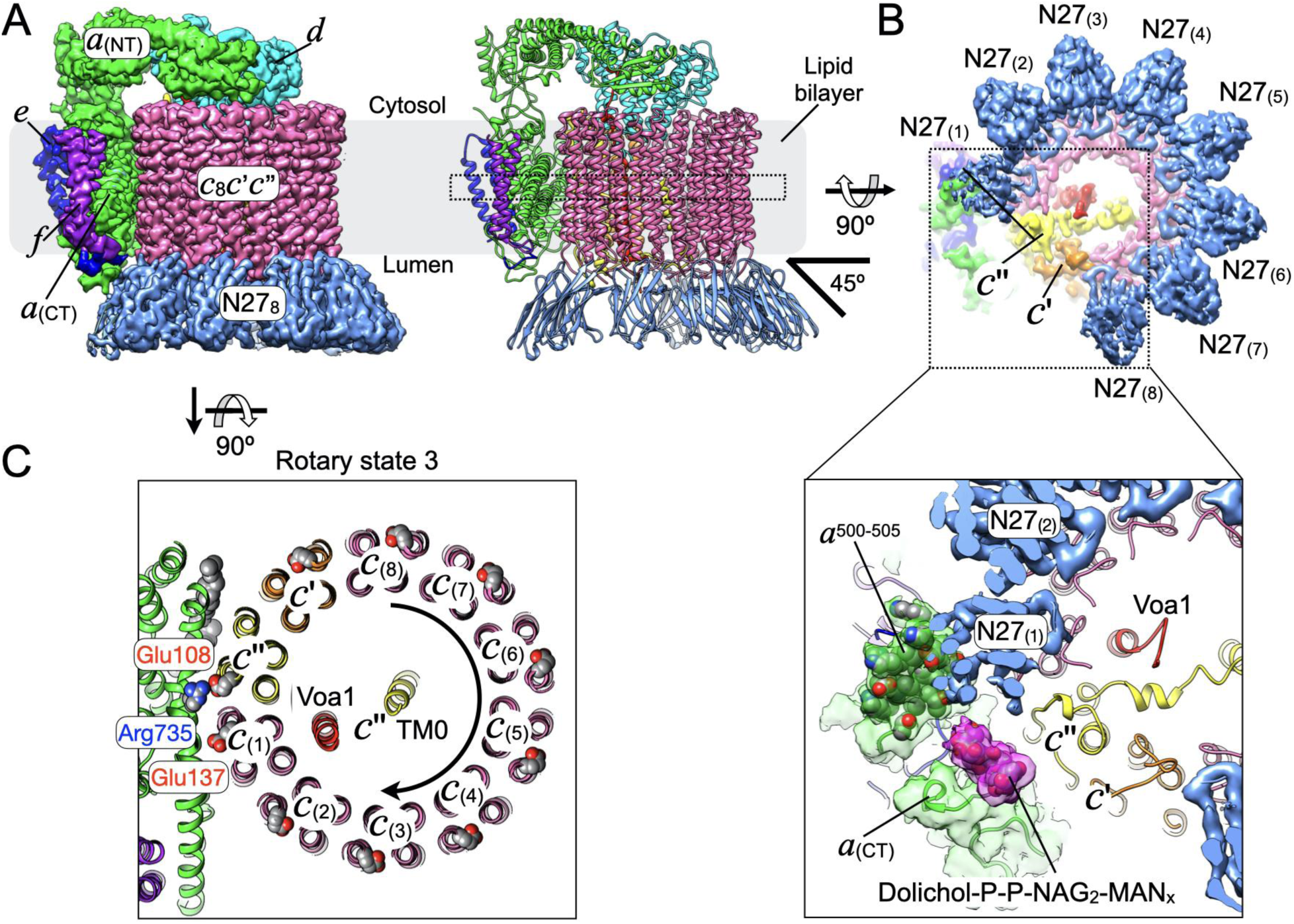
N27 binds the luminal side of the *c*-ring. **A** Left: 3.3 Å resolution EM density of V_o_ND bound to eight copies of Nb N27, showing density for subunits *a* (green), *d* (cyan), *e* (blue), *f* (purple), and N27 Nbs (blue). Right: Ribbon representation, showing that N27 Nbs bind the *c*-ring at an angle of ∼45°. **B** Upper panel: View from the lumen towards the membrane showing the EM density for the eight N27 Nbs bound to the eight *c* subunits of the *c*-ring. Note the reduced density for N27_(1)_ and N27_(8)_. Lower panel: Potential steric clash (hindrance) between N27 Nbs and *a*_CT_ loop residues 500-505 and the glycolipid ligand (Dolichol-P-P-NAG2-MANx), which is an integral component of the V_o_ (Roh et al, 2020). Coordinates for the dolichol ligand are from 6m0r (Roh et al, 2020). **C** Cross section at the level of the conserved glutamic acid and arginine residues in the proteolipid subunits (*c*, *c’*, *c”*) and subunit *a*, respectively. In the canonical rotary state 3 of the isolated V_o_, the glutamic acid of *c*’’ (Glu108) is in contact with Arg735 of *a*_CT_.

**Figure 6:**
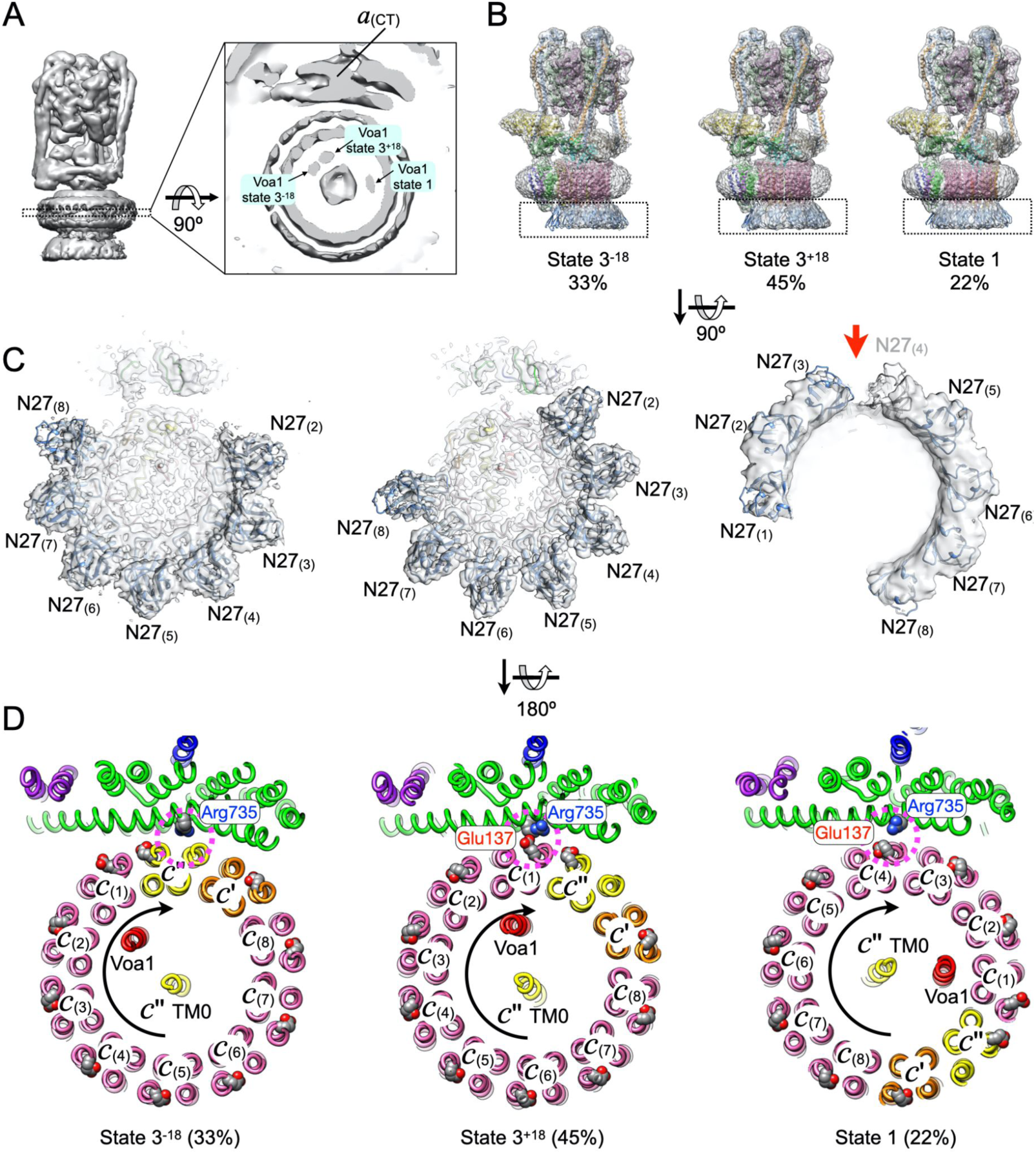
N27 binds the luminal side of the *c*-ring. **A** Low-resolution reconstruction of N27 bound V_1_(H_chim_)V_o_ND showing presence of three rotary states as evident from the position of the density for Voa1. **B** The three rotary states in (A) were separated by 3-D classification into states referred to as 3^-18^, 3^+18^, and the canonical state 1. **C** View from the lumen towards the N27 Nbs. Note the lack of density for N27_(1)_ in states 3^-18^ and 3^+18^. A gap in density for the state 1 complex near *a*_CT_’s loop and the glycolipid indicates low occupancy and/or increased dynamics for N27_(3)_ and N27_(4)_ (see red arrow). Note that the state 1 map was low pass filtered for representation. **D** Cross section at the level of the conserved glutamic acid (red spheres) and arginine residues (blue spheres) in the proteolipids and subunit *a*, respectively. In rotary state 3^+18^ (middle panel), Arg735 is in contact with Glu137 of *c*_(1)_ (see pink dashed circle). In rotary state 1 (right panel), Arg735 is close to Glu137 of *c*_(4)_ (see pink dashed circle). Note that in state 3^-18^, none of the *c*-ring glutamates appears to be close enough to form a salt bridge with Arg735 of *a*_CT_.

### Structure of N27 bound V_o_ND subcomplex

The map of N27 bound V_o_ND shows that this Nb binds the eight proteolipid *c* subunits (Vma3p) on the luminal side of the complex (**Fig. 5A**), but not the proteolipid isoforms *c*’ (Vma11p) and *c*’’ (Vma16p) (**Fig. 5B**, upper panel). N27 is bound with the long axis of the Nb at an angle of ∼45° with respect to the long axis of the *c*-ring, giving the N27-bound V_o_ND the appearance of an inverted crown (**Fig. 5A**). The N27 bound to the *c* subunit adjacent to *c*’’ (*c*_(1)_; N27_(1)_) makes contact with a luminal loop of *a*_CT_ formed by residues 500-505 (**Fig. 5B**, lower panel). Rotation of the N27 bound *c*-ring (which is part of the rotor) would result in a steric clash between the Nb and the luminal loop of *a*_CT_ (which is part of the stator), a clash that is likely responsible for N27’s ability to inhibit the holoenzyme’s ATPase activity. Another clash during *c*-ring rotation would likely occur between N27s and the glycolipid (dolichol-P-P-NAG_2_-MAN_x_), which is an integral part of yeast *a*_CT_ (Roh et al, 2020) (**Fig. 5B**, lower panel). An analysis of the N27:subunit *c* interface using the PISA server (Krissinel & Henrick, 2007) revealed that most of the contacts on the *c* subunit’s side are mediated by its N- and C-terminal residues, as well as residues contained in the luminal loop connecting transmembrane ⍺ helices (TMHs) 2 and 3 (**Appendix Fig. S7A**). Contacts are also observed between individual Nbs (**Appendix Fig. S7B**), which may contribute to the sub-picomolar binding affinity (avidity) of N27 as measured using BLI (**Fig. 4**).

### Structure of N27 bound holoenzyme

The N27 bound V_o_ND is locked in the canonical rotary state 3, in which the conserved glutamic acid residue of *c*’’ (Glu108 in yeast) is engaged in a salt bridge with the conserved arginine residue in *a*_CT_ (Arg735) (**Fig. 5C**). In our recent cryoEM structural analysis of the *in vitro* reconstituted yeast V-ATPase containing V_1_H_chim_, we obtained maps for the three rotary states with 59% of the particles in state 1, 30% in state 2 and 11% in state 3 (Khan et al, 2022), similar to what had been described earlier for the detergent solubilized wild type enzyme (Zhao et al, 2015). Since *in vitro* reconstitution starts from autoinhibited V_o_ND that is exclusively locked in state 3, we reasoned that once assembly with V_1_H_chim_ and subunit C has occurred, thermal energy alone is sufficient (even in absence of added nucleotides) to overcome energy barriers and establish the observed distribution of rotary states, given enough time and depending on temperature (Khan et al, 2022). As mentioned above, the cryoSPARC 3-D reconstruction of N27 bound holoenzymes revealed significant conformational heterogeneity as reflected in poor density for the V_o_ region including bound Nbs compared to the map of N27 bound V_o_ND (**Appendix Fig. S4**). We therefore analyzed the dataset in more detail. A low resolution 3-D reconstruction of all N27-bound holoenzyme particles revealed three positions of the density for Voa1p within the cavity of the *c*-ring (**Fig. 6A**). Subsequent 3-D classification and refinement resulted in maps for rotary state 1 (22%) and two state 3-like conformations referred to as states 3^-18^ and 3^+18^ (together 78%) (**Fig. 6B**; **Appendix S6**). To facilitate interpretation of the N27 bound V_o_ regions, we performed focused refinement of three different rotary states. Maps for the V_o_ subcomplexes for states 3^-18^ and 3^+18^ show density for seven N27 Nbs, with essentially no density for N27_(1)_. The state 1 map is not as well resolved but has clear density for seven N27 Nbs, with N27_(4)_ only partially resolved due to a gap in density adjacent to the luminal loop of subunit *a*. (**Fig. 6C**, see red arrow). At a lower contour level, however, some density is visible for for N27_(4)_, indicating lower Nb occupancy or increased dynamics at this site. In rotary state 1, the conserved glutamic acid residue of *c*_(4)_ is engaged in a salt bridge with conserved arginine 735 in *a*_CT_ (**Fig. 6D**, right panel). The observed rotary state 3^+18^ of the N27 bound holoenzyme (∼45%) has so far only been observed for the isolated, autoinhibited V_o_ (Roh et al, 2020), but not V_1_V_o_. Compared to the canonical rotary state 3, the *c*-ring in the state 3^+18^ configuration is rotated ∼18° clockwise towards state 1. In this configuration, Arg735 of *a*_CT_ is in contact with Glu137 of the *c* subunit adjacent to *c*’’ (*c*_(1)_) (**Fig. 6D**, middle panel) instead of Glu108 of *c*’’ as observed in the canonical rotary state 3 (**Fig. 5C, Appendix S8**). In the third class of N27 bound holo V_1_(H_chim_)V_o_ND (33%), the *c*-ring is rotated ∼18° counterclockwise towards state 2 (**Fig. 6D**, left panel). In this hitherto unseen rotary sub-state referred to as 3^-18^, however, none of the *c*- ring glutamates seem to be engaged with *a*_CT_’s Arg735. Overall, the analysis shows that N27 binding to *in vitro* reconstituted V_1_(H_chim_)V_o_ND stabilizes rotary states not seen previously, conformations that may ultimately provide insight into the mechanism of V-ATPase assembly.

### Epitope of N125

CryoEM analysis of V_o_ND complexes bound to N125 revealed that N125 (much like N27) binds the *c*-ring on the luminal side of the complex. 3-D classification revealed that ∼76% of the particles have four, and the remaining ∼24% three copies of N125 bound (**Fig. 7A**; **Appendix Fig. S9**). We refined the best resolved class of particles having four N125 bound to a final resolution of 3.4 Å (**Fig. 7B**). The map shows that N125 occupies a primary *c* subunit, but also some surface area of a neighboring *c* subunit, which results in N125 binding to alternating *c* subunits, for a maximum occupancy of four Nbs (**Fig. 7C,D**). Due to N125 binding with the long axis of the Nb aligned with the long axis of the *c*-ring, there appears to be no significant steric clash with the luminal loops of *a*_CT_ upon *c*-ring rotation, consistent with this Nb having no significant effect on the holoenzyme’s ATPase activity (**Fig. 3A**). CryoEM analysis of a dataset of ∼24,000 N125 bound V_1_(H_chim_)V_o_ND particles revealed that N125 binds to the intact enzyme in a similar fashion compared to V_o_ND (**Fig. 7E**). The limited size of the dataset, however, did not permit sorting of the particles based on rotary states and Nb occupancy.

**Figure 7:**
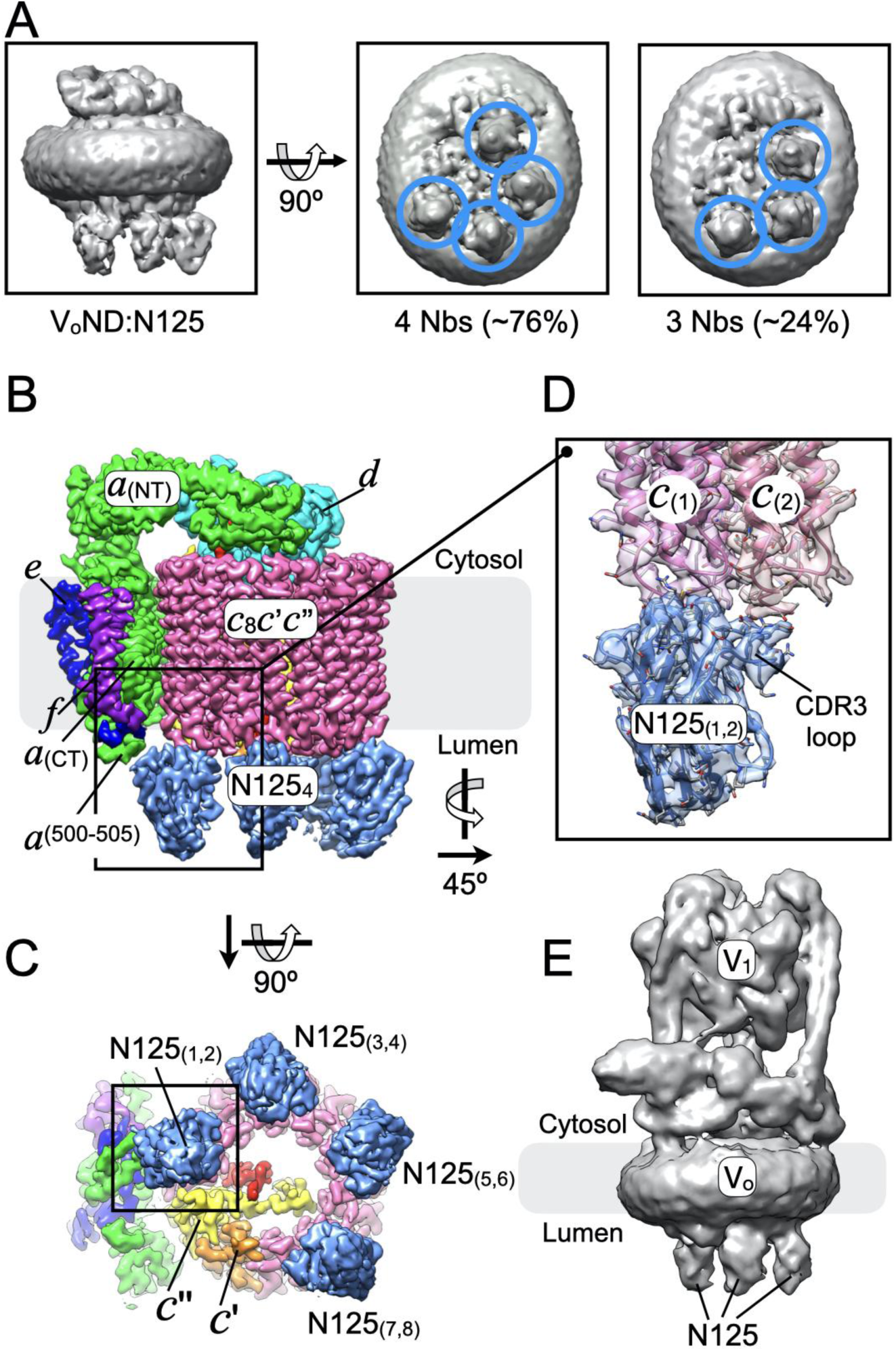
N125 binds the luminal side of the *c*-ring. **A** Low-resolution reconstruction of N125 bound V_o_ND. 3-D classification reveals classes with either three (middle panel) or four (right panel) Nbs bound. Nbs are highlighted by the blue circles. **B** 3.4 Å reconstruction of V_o_ND bound to four N125 Nbs. As N125 binds with the long axis parallel to the long axis of the *c*-ring, there is no significant contact between the Nbs and *a*_CT_. **C** The N125 Nbs occupy two *c* subunits each for a maximum ratio of 4 Nbs per complex. **D** N125_(1,2)_ binds primarily to *c*_(1)_ but the Nb’s CDR3 loop makes a significant contact with *c*_(2)_. **E** Low-resolution map of N125 Nb bound to V_1_(H_chim_)V_o_ND obtained from a dataset of ∼24,000 particles.

### N125 inhibits activity when tethered to NeutrAvidin

Since N125 binding does not result in significant inhibition of V_1_(H_chim_)V_o_ND’s ATPase activity (**Fig. 3A**), we wished to determine if adding a large globular protein to the Nb’s C-terminus could create an inhibitory effect due to steric hindrance of *c*-ring rotation. To do so, we expressed N125 with a cleavable C-terminal Biotin Acceptor Protein (BAP)-tag (N125^HBP^). We then measured ATPase activity of N125^HBP^ bound V_1_(H_chim_)V_o_ND in presence of tetrameric NeutrAvidin (∼60 kDa). The data show that NeutrAvidin-tethered N125^HBP^ reduced V_1_(H_chim_)V_o_ND’s ATPase activity to ∼42% of the control, with NeutrAvidin alone having no significant effect on the activity (**Fig. 8**). Thus, addition of a ∼60 kDa globular protein to the C-terminus of N125 converts the non-inhibitory Nb to an inhibitory one.

**Figure 8:**
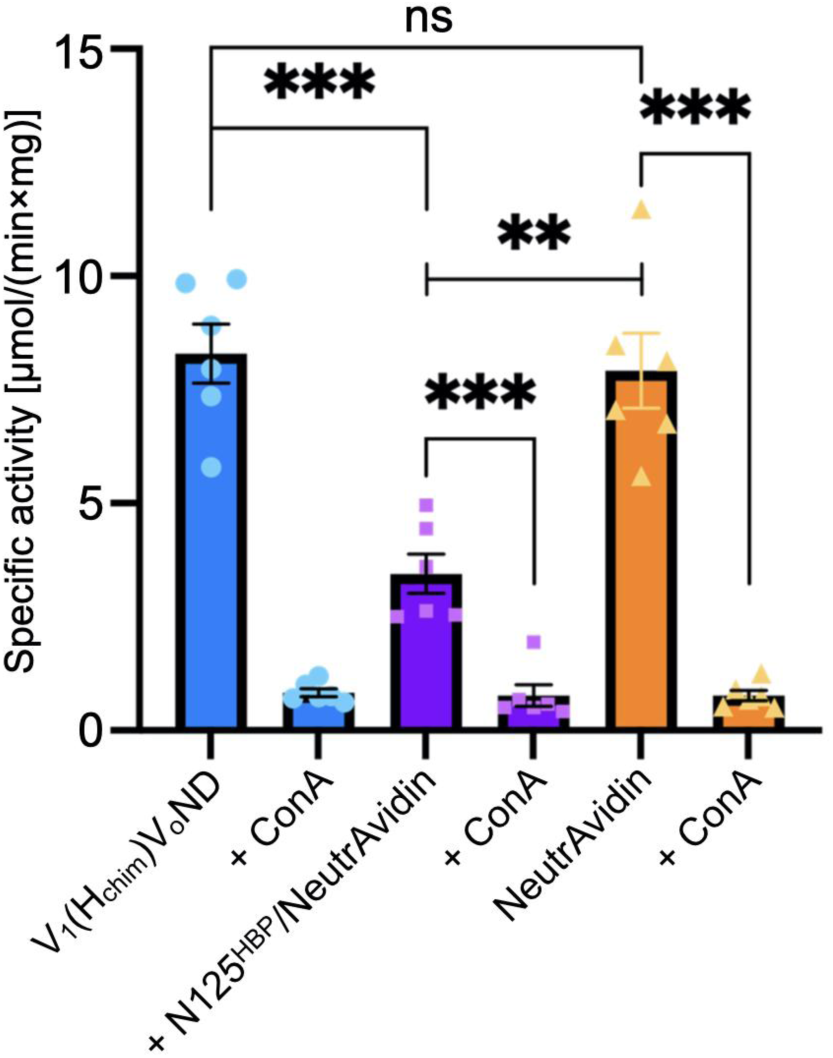
NeutrAvidin bound N125 activity inhibition. The specific activity of V_1_(H_chim_)V_o_ND is on average 8.30 µmol/(min×mg), and when inhibited by ConA is reduced to 0.84 µmol/(min×mg) (blue). When V_1_(H_chim_)V_o_ND is incubated with N125^HBP^-NeutrAvidin, specific activity is reduced to 3.45 µmol/(min*mg) prior to additional inhibition by ConA (0.77 µmol/(min×mg)) (purple). NeutrAvidin alone does not inhibit V-ATPase activity (orange). Measurements are three technical replicates each from two independent purifications of V-ATPase and two independent purifications of Nbs. Error bars are SEM. p values were calculated using unpaired t test assuming different SD in GraphPad Prism software, with ns>0.05, **≤0.01, and ***≤0.001. The specific V-ATPase inhibitor, ConcanamycinA, ConA was used as a control.

### Epitope of N2149

The activity and assembly assays performed with N2149 (**Fig. 3**) showed that while the Nb had no significant effect on the activity of the assembled complex, binding to V_o_ND prior to assembly prevented binding of V_1_H_chim_. This suggested that N2149 binds the cytosolic side of the complex and that the epitope was located at or near one of the V_1_ binding sites. Analysis of the N2149:V_o_ND complex using cryoEM resulted in a 3.7 Å map that showed the Nb bound to subunit *d* (**Fig. 9A; Appendix Fig. S10**). Here, the CDR3 loop of the Nb is seen to partially insert into the central cavity of *d*, which serves to bind V_1_ subunit D in the assembled complex (**Fig. 9B**). Indeed, overlaying the structure of N2149-bound subunit *d* with that of unbound subunit *d* as part of state 3 holoenzyme (PDB 7FDC) revealed that N2149’s CDR3 loop would clash with subunit D and prevent proper V_1_ binding, consistent with N2149 inhibiting the assembly of V_1_H_chim_ with V_o_ND (**Fig. 3** and **Fig. 9C**, right panel).

**Figure 9:**
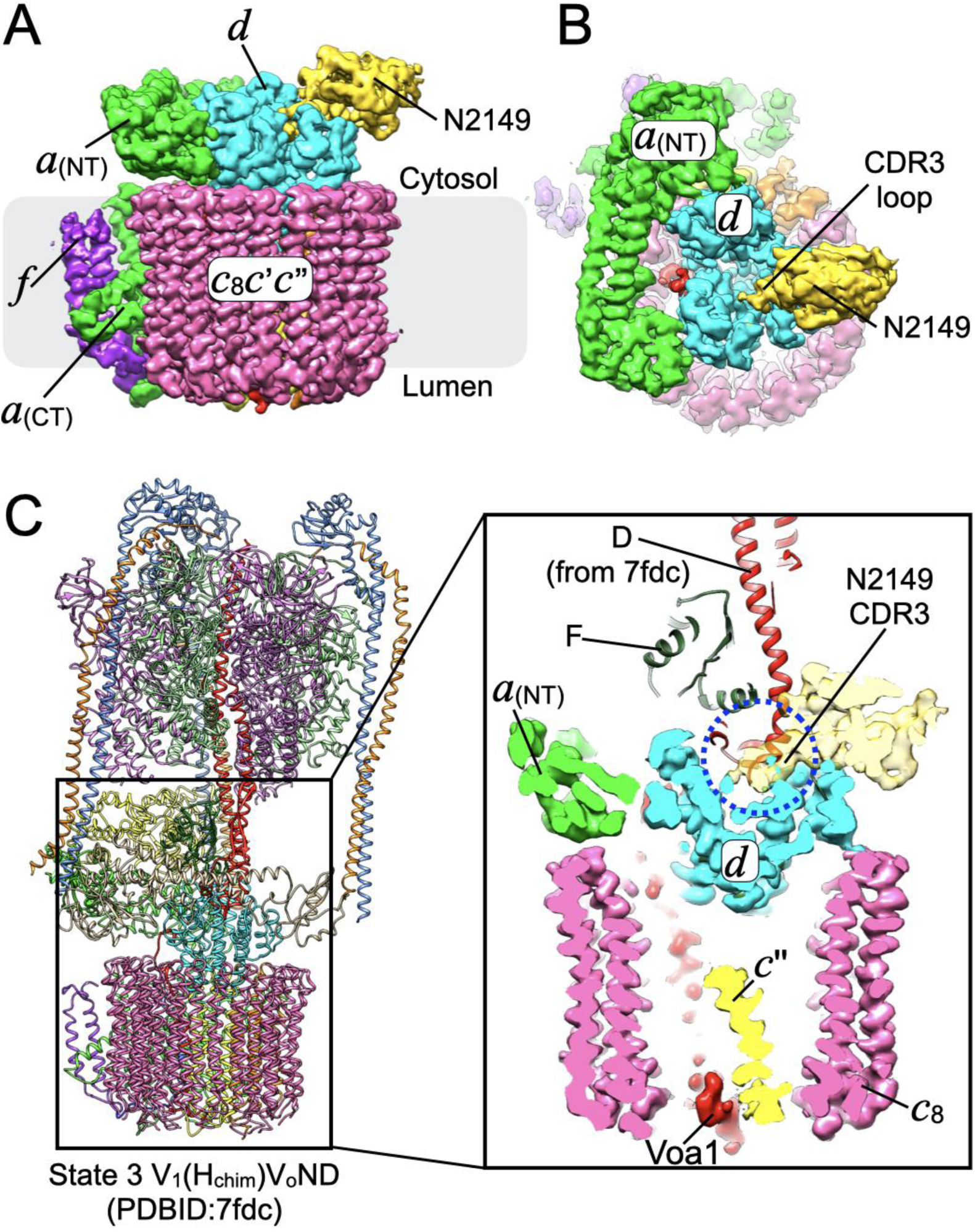
N2149 binds to subunit *d*. **A** N2149 (gold) binds to subunit *d* (cyan) in the V_o_ND subcomplex. **B** Top-view of N2149-bound V_o_ND, showing that the CDR3 loop of the Nb inserts into the central cavity of the *d* subunit. **C** Left panel, state 3 V_1_(H_chim_)V_o_ND (PDBID:7fdc). Right panel, the N2149 bound subunit *d* from the V_o_ND complex shown in panel (A) is overlaid on subunit *d* as part of the state 3 holoenzyme, indicating that the CDR3 loop of the Nb would occupy the binding site of the V_1_-D subunit in the intact complex.

### N125 immunoprecipitates human V-ATPase

Eukaryotic V-ATPase is highly conserved on the structural and primary sequence level from yeast to mammals, including humans. Determining the ability of Nbs to bind to and regulate activity of the yeast V-ATPase was a critical first step, but identifying Nbs that have affinity for the human VATPase will be an essential prerequisite for developing Nbs as therapeutics. A recent study in mice showed that inhibition of the plasma membrane localized V-ATPase with a Nb that binds at the extracellular side of the *c* subunit inhibited lung metastasis from grafted tumors (Li et al, 2024). We therefore wanted to test if any of the ⍺-yeast V_o_ND Nbs bind the human V-ATPase (HsV_1_V_o_). Since our N125^HBP^ construct permits biotinylation of the Nb for use in streptavidin-based pull-down with low background, we chose to test whether N125 can immunoprecipitate purified human V-ATPase. Streptavidin (SA) beads were coated with purified, biotinylated N125^HBP^ and incubated with a mixture of lipid nanodisc reconstituted purified human holo V-ATPase (*Hs*V_1_V_o_ND) and proton channel subcomplex (*Hs*V_o_ND). N125- bound protein was “eluted” from the beads using HRV-3C protease (also referred to as Prescission protease) to cleave the biotinylated BAP-tag on the Nb (**Figure 10A**). Silver-stained SDS-PAGE gels of eluate fractions revealed presence of both V_1_ and V_o_ subunits, indicating that the ⍺-yeast N125 Nb was able to pull down the human V-ATPase under these conditions (**Fig. 10B**). Attempts to visualize N125 bound to the human V-ATPase using cryoEM, however, were not successful, presumably due to the reduced affinity of the ⍺-yeast Nb for the human enzyme. Affinity of N125 for the human V_o_ could be reduced due to limited sequence conservation for the N- and C-termini and the luminal loop of yeast and human *c* subunits **(Appendix Figure S11)**. Moreover, the human V_o_ contains Ac45 (ATP6AP1), which forms a heavily glycosylated “plug”-like structure on the *c*-ring cavity exposed to the organelle lumen or the extracellular side and which could limit access of the Nbs to the *c*-ring.

**Figure 10:**
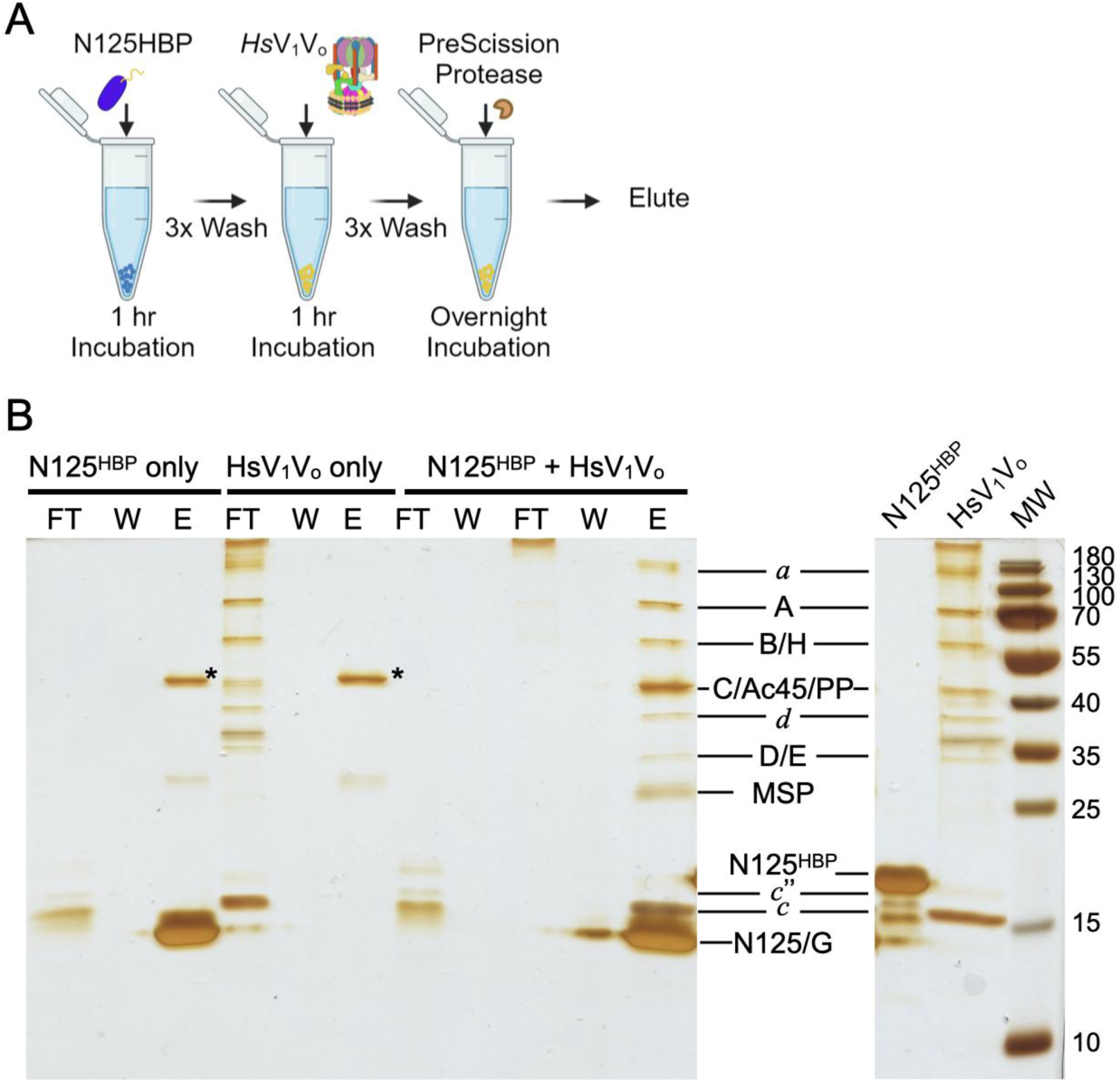
N125HBP Pulldown of *Hs*V_1_V_o_. **A** N125^HBP^ (containing a biotin tag) was bound to Streptavidin (SA) beads, washed and incubated with (or without) *Hs*V_1_V_o_ before an overnight elution with PreScission Protease (PP) to release N125 from the beads. *Hs*V_1_V_o_ was also applied to empty SA beads (*Hs*V_1_V_o_ only) as a non-specific binding control. **B** Silver stained 15% SDS-PAGE of pull-down samples and controls. From left to right: N125^HBP^ only control (no *Hs*V_1_V_o_), *Hs*V_1_V_o_ only control (no Nb on beads), N125^HBP^ + *Hs*V_1_V_o_; FT = flow through, W = wash, E = elution by release of N125 with protease. The band corresponding to Prescission protease in the control elution samples is indicated by the asterisk (*). The right panel shows purified “input” samples N125^HBP^ and HsV_1_V_o_.

## Discussion

The V-ATPase is an essential enzyme in all cells of higher eukaryotic organisms as most, if not all, cellular functions depend on differential pH across cellular membranes (Collins & Forgac, 2020; Eaton et al, 2021; Forgac, 2007; Futai et al, 2019). Not surprisingly, aberrant V-ATPase activity is linked to numerous human diseases, and while most are due to loss of enzyme function (such as renal acidosis, hearing loss, neurodegeneration) (Beauregard-Lacroix et al, 2021; Guerrini et al, 2022; Karet, 1999; Smith et al, 2000), there are cases where the disease state is linked to normal V-ATPase activity (such as viral infection) (Lindstrom et al, 2018; Marjuki et al, 2011). Furthermore, disease states have also been linked to normal activity at the wrong place due to enzyme mislocalization. Indeed, highly invasive and metastatic tumor cells display V-ATPases at the plasma membrane in cell types that do not normally display that pattern of localization (Kulshrestha et al, 2015; Sennoune, 2004; Stransky et al, 2016). While V-ATPase has been recognized as a drug target (Kartner, 2014; Niikura, 2006), there are challenges with developing therapeutic agents. First, V-ATPases are found on virtually every cell membrane, and total loss of enzyme function is lethal to all mammalian cells (Inoue et al, 1999), thus excluding application of pan V-ATPase inhibitors. Second, most mammalian V-ATPase subunits are expressed as multiple isoforms, some of which are tissue and/or cell location specific (Toei et al, 2010). One first step in targeting V-ATPase will therefore be to develop therapeutics that can specifically target a disease-causing V-ATPase population without inhibiting populations that are essential for cell function and survival. Recently, studies have shown that targeting mislocalized V-ATPases on the plasma membrane of tumor cells using monoclonal antibodies (in case of ovarian cancer) or Nbs (in case of xenografted tumors in mice) can induce tumor cell apoptosis and reduce lung metastases, respectively (Kulshrestha et al, 2020; Li et al, 2024).

Nbs are particularly promising for therapeutic use due to their stability, small size, and ease of production. With diffusion being the primary mode of transportation within tumor sites due to low lymphatic drainage, the small size, and therefore faster diffusion rate, of Nbs allows them to achieve a homologous distribution more quickly than traditional antibodies (Thurber et al, 2008). Additionally, the monovalent nature of Nbs allows them to avoid the so-called “site barrier effect” during which bivalent therapeutics get trapped on the periphery of tumor sites (Chames et al, 2009; Xenaki et al, 2017). However, the rapid rate of renal clearance, which prevents accumulation of the therapeutic at the tumor site, is a major drawback of Nb use, and it has been proposed that treatment using Nbs may need to occur with greater frequency to overcome this clearance effect (Jin et al, 2023). The rapid clearance of Nbs from the body does not exclude its use in therapeutics, and the small size and increased specificity of Nbs is useful for treating a variety of diseases. Therefore, several Nbs have been evaluated in clinical trials for their potential use in treatment of autoimmune disease, lymphoma as well as brain and breast cancer (Jovcevska & Muyldermans, 2020; Muyldermans, 2021). Nbs are particularly useful for targeting and treating infectious diseases as Nbs can be generated against conserved conformational epitopes that are inaccessible to traditional antibodies, and with the comparatively fast generation of Nbs new therapeutics can be made to keep up with the rapid mutations viruses are known to undergo. For these reasons, there are numerous Nbs in clinical trials for the treatment of infectious diseases such as SARS-CoV-2, HIV, Hepatitis B, and Influenza Virus (Alexander & Leong, 2024).

Here we have generated and characterized Nbs raised against the yeast V-ATPase V_o_ proton channel complex. Nbs were generated at the Vrije Universiteit Brussel’s VIB Nanobody facility, resulting in 94 Nb clones. For llama immunization, phage display and ELISA based Nb selection, the purified yeast V_o_ complex was reconstituted into lipid nanodiscs (V_o_ND).

Previously, we obtained a 2.7 Å cryoEM structure of V_o_ND (Roh et al, 2020), and we showed that lipid nanodisc reconstitution of holo V-ATPase results in superior stability compared to the detergent solubilized enzyme (Sharma et al, 2019; Sharma & Wilkens, 2017; Uchida et al, 1988). Small scale high-throughput screening showed that most of the Nbs not containing internal Amber or Opal stop codons could be expressed and purified from *E. coli*. We then tested a subset of the Nbs from different CDR groups and based on the ELISA data provided by the VIB Nb facility, we chose three Nbs for a more detailed biochemical characterization. From the ATPase activity and assembly assays, we determined that each of the three Nbs tested showed a unique behavior toward the V-ATPase, with one Nb (N27) resulting in almost complete inhibition of the activity, a second one (N2149) preventing assembly of the holoenzyme from purified components *in vitro*, and a third one (N125) having little to no impact on activity or assembly. This diversity of effects on V-ATPase activity and assembly from only a small sampling of our Nb collection reflects the benefits of using the V_o_ subcomplex for immunization, rather than a single subunit.

The Nbs impact on activity or assembly can be rationalized by their binding epitopes, which we determined using cryoEM structure determination. The inhibitory N27 binds the luminal side of the *c* subunits at an angle of ∼45°, which allows the Nb to occupy a maximum of all eight subunits of the *c*-ring (as observed for the V_o_ND:N27_8_ complex). On the other hand, the ∼45° binding is expected to cause friction with *a*_CT_ and the integral glycolipid during *c*-ring rotation. BLI based binding kinetics indicated that N27 binds V_o_ND with a sub-picomolar K_d_ due to virtually no significant dissociation over the time period of the experiment. Analysis of the Nb binding interactions using the PISA server, however, revealed that N27 shares buried surface area not only with the *c* subunit (∼530 Å^2^ for N27_(8)_:*c*_(8)_), but also with neighboring N27 Nbs (∼240 Å^2^ for the N27_(8)_:N27_(7)_ interface), Nb-Nb interactions that may result in increased binding avidity. Support for this assumption comes from the observation that the N27s which have only one neighbor display reduced density in the cryoEM map of the N27:V_o_ND complex.

Surprisingly, cryoEM of the N27 bound V_1_(H_chim_)V_o_ND complex did not produce a class of complexes halted in the canonical rotary state 3. Instead, the analysis revealed two state 3-like sub-states, which we designated 3^-18^ and 3^+18^ based on the rotational deviation from state 3.

Whereas the 3^+18^ sub-state had been described before for V_o_ND, the 3^-18^ sub-state has not been seen previously and while the resolution of the map does not allow accurate placement of side chains, the fitted structure suggests that there is no contact between *a*_CT_’s Arg735 and any of the conserved *c*-ring glutamic acid residues. The reason why the canonical state 3 is observed for isolated N27 bound V_o_ND but not for the N27 bound holoenzyme is not clear at the moment, but it is possible that the 3^+18^ and 3^-18^ sub-states are early assembly intermediates stabilized by N27 binding.

The straight binding mode of N125 is likely the reason why this Nb does not inhibit *c*-ring rotation. N125, however, becomes inhibitory when tethering its C-terminus to a ∼60 kDa globular protein. Despite the lack of significant sequence conservation between yeast and human *c* subunit N- and C-termini, N125 was able to pull down human V-ATPase, suggesting that part of the binding energy is provided by shape complementation in addition to specific residue contacts. The affinity for the human enzyme, however, is significantly lower as we were not able to observe N125 bound to the human V_o_ using cryoEM. It is possible that binding of N125 to the human *c*-ring is facilitated by the high local concentration of the Nbs on the streptavidin beads due to avidity effects, but once Nbs are released from the beads, individual Nbs will dissociate due to the loss of avidity effects on the beads. The finding, however, that an ⍺-yeast Vo Nb can bind the human V_o_ is promising and suggests the possibility that N125 (or any of the other *c*-ring binding Nbs) can be engineered for improved binding and inhibition. Work by the Forgac laboratory showed that *a*3 and *a*4 containing V-ATPases are mislocalized to the plasma membrane in cells derived from highly aggressive breast tumors, and that inhibition of the mislocalized complexes from the outside of the cell via antibody binding to an engineered affinity tag at the *c* subunit C-terminus reduced cell invasiveness and migration (Capecci & Forgac, 2013; Cotter et al, 2015; McGuire et al, 2019; Nishisho et al, 2011). A follow-up study in mice showed that administration of a bifunctional Nb against the mouse V-ATPase *c* subunit N-terminus selected from a naive Nb library suppresses lung metastasis from xenografted tumors (Li et al, 2024). Thus, identifying Nbs that inhibit the human enzyme by binding to the lumen side of the complex will be a first step in developing Nbs as a V-ATPase based anti-cancer therapy.

In contrast to the other two Nbs, we found that N2149 binds on the cytosolic side of the complex to rotary subunit *d*, a single copy subunit that acts as an adapter linking V_1_ subunit D to the V_o_ complex. N2149 binds in a position that would block V_1_ binding to V_o_, in accord with our functional analyses. Indeed, N2149 inhibits assembly of V_1_H_chim_ and V_o_ND *in vitro* but not V-ATPase activity, suggesting an inability to bind holoenzymes. V-ATPase is regulated by reversible disassembly of the complex into isolated V_1_ and V_o_ subcomplexes. Our data indicates that N2149 binds to an epitope that is only revealed when the complex is disassembled. A traditional antibody raised against *a*_NT_ that recognizes a cryptic epitope only available in V_o_ has been a widely used workhorse for the study of reversible disassembly via immunoprecipitation and western blotting (Kane, 1995). Since Nbs can be both easily genetically modified (eg. fluorescent tag) and expressed in cells, N2149 may represent a useful tool for studying V-ATPase disassembly in cells.

In conclusion, Nbs show great promise in use as tools to identify disassembled V-ATPase, modulate V-ATPase activity, or be used to purify endogenous V-ATPase populations. Lastly, some Nbs have the potential to be developed into cancer therapeutics aimed at inhibition of metastasis. While more research is still needed in this area, our work highlights the potential of Nbs in targeting the V-ATPase for use in functional studies or the development of therapeutics.

## Materials and Methods

### Subunit C and H_chim_ expression and purification

Subunit C and H_chim_ were expressed and purified as described previously (Oot & Wilkens, 2010; Sharma et al, 2019). Briefly, both proteins were expressed in Rosetta 2 *E. coli* as N-terminal fusions with a cleavable (via HRV-3C, PPase) maltose-binding protein (MBP) tag. Cells were grown to an OD_600_ 0.6 in rich broth (LB + 2% glucose) supplemented with 50 µg/ml carbenicillin and 34 µg/ ml chloramphenicol. Expression was induced with 0.5 mM IPTG at 30 °C and cells were allowed to grow for another 5.5 hours. Cells were harvested by centrifugation at 3,500 ⨯ g, resuspended in amylose column buffer (ACB) (20 mM Tris-HCl, 200 mM NaCl, 1 mM EDTA, pH 8), and stored at -20 °C. Cells were thawed, treated with lysozyme and DNAse, sonicated, and centrifuged at 13,000 x g for 30 min at 4 °C. The cleared supernatant was passed over a 20 ml amylose column, pre-equilibrated with 10 CV ACB, washed with 10 CV ACB + 1 mM DTT, and eluted with 25 ml ACB containing 10 mM maltose and 1mM DTT. The MBP tag was cleaved using PPase for 2 hours in presence of 5 mM DTT. After cleavage, ion exchange chromatography was used to separate MBP from subunit C, or H_chim_ as follows. The cleavage product containing subunit C was dialyzed into 20 mM Bis-Tris-HCl pH 6.5, and 0.5 mM EDTA before application to a diethylaminoethyl (DEAE) column, with MBP binding to the anion exchange resin and C subunit passing through the column. The cleavage product containing H_Chim_ was dialyzed into 25 mM sodium phosphate, pH 7, 5 mM β-ME, and 0.5 mM EDTA before loading onto a carboxymethyl (CM) column, where Hchim binds to the cation exchange resin and MBP flows through the column. Both subunit C and, H_Chim_ were purified further by size exclusion chromatography using Superdex 75 for subunit C and Superdex200 column for H_chim_ attached to an Äkta FPLC and stored in 20% glycerol at -80 °C.

### Expression and purification of membrane scaffold protein (MSP) for nanodisc (ND) reconstitution

MSP was expressed and purified as previously outlined in (Oot & Wilkens, 2024; Oot et al, 2021) with modifications. Briefly, *E. coli* cells (BL21DE3) harboring a pET28a plasmid encoding an N-terminal 6xHis-BAP-PPase-tagged MSP1E3D1 were grown in rich broth at 37 °C. Media was supplemented with 50 µg/ml kanamycin before induction at OD_600_ 0.5-0.6 with 0.5 mM IPTG. Cells were grown for 4 hours at 37 °C post induction before being harvested by centrifugation at 3,000 rpm for 25 minutes. Pellets were resuspended in lysis buffer (20 mM Tris-HCl, 500 mM NaCl, 10 mM Imidazole, 6 M Guanidine-HCL, pH 8), and pelleted again at 3,000 rpm for 15 minutes before discarding the supernatant and storing at -20 °C. Cells were gently resuspended in 5 ml lysis buffer per gram of cell pellet. Cells were lysed by sonication, 2 x 20 seconds on ice. Broken cells were centrifuged at 13,000 x g for 40 min at room temperature. Supernatant containing MSP was purified over a 10 ml Ni-NTA column at room temperature, pre-equilibrated with 5 CV lysis buffer. The column was washed with 5 CV of lysis buffer, followed by 10 CV Buffer A (20 mM Tris-HCl, 10 mM imidazole, 250 mM NaCl, pH 8). The following step was added when purifying MSP to be used for reconstitution of HsV_1_V_o_ into nanodiscs: the column was then washed with 6 CV Buffer B (20 mM Tris-HCl, 300 mM NaCl, 1% TritonX-100, pH 8), 10 CV buffer C (20 mM Tris-HCl, 300 mM NaCl, 25 mM sodium cholate, pH 8), and finally 10 CV of buffer A. MSP was eluted from the column with 5 CV buffer B (20 mM Tris-HCl, 1 M Imidazole, 250 mM NaCl, pH 8) and fractions analyzed using SDS-PAGE. Peak fractions were pooled, and in case of MSP used for nanodisc reconstitution of yeast Vo, the His-BAP tag was cleaved using PPase and dialyzed overnight into 20 mM Tris-HCl, 100 mM NaCl, 1 mM DTT before re-applying to an Ni-NTA column to remove the cleaved tag.

Cleaved MSP was concentrated using 20 ml VivaSpin 5000, protein concentration was determined using UV absorbance at 280 nm. In the case of MSP to be used for HsV_1_V_o_ nanodisc reconstitution, purified MSP was dialyzed into 4 mM Tris-HCl, 15 mM NaCl, pH 8, divided into 60 mg batches and lyophilized for storage at -80 °C.

### V_1_ΔH expression, purification, and V_1_H_Chim_ reconstitution

V_1_ΔH was expressed and purified as described (Sharma et al, 2019). Briefly, *Saccharomyces cerevisiae* harboring genomic deletions of both subunits H (VMA13) and G (VMA10) (*vma13Δ::KanMX*, *vma10Δ::Nat*) (Oot, 2016; Zhang et al, 2003) was transformed with a pRS315 (*Leu*) plasmid carrying an N-terminally FLAG tagged subunit G and maintained in SD-Leucine (SD -Leu) medium to maintain the plasmid. Twelve liters of cells were grown at 30 °C overnight at 200 rpm and harvested at OD^600^ 3.0-4.0 by centrifugation at 3,000 rpm for 25 minutes at 4 °C. Cell pellets were resuspended in TBSE (20 mM Tris-HCl, 150 mM NaCl, 0.5 mM EDTA, pH 7.2) and stored at -80 °C in 6 l portions until use. For purification, cells from 6 l culture were brought to 100 ml in TBSE and supplemented with 5 mM β-ME and protease inhibitors before lysis using 16-20 passes through a microfluidizer, with 5 min breaks every 3 or 4 passes to prevent overheating. Lysed cells were centrifuged at 4,000 rpm for 30 min at 4 °C, and the supernatant then centrifuged at 13,000 ⨯ *g* for 1 h at 4 °C to remove unbroken cells and mitochondria, respectively. The supernatant was applied to a pre-equilibrated 5 ml ⍺-FLAG column pre-equilibrated with 10 CV TBSE, and eluted with 5 CV of TBSE containing 0.1 mg/ml FLAG peptide. Fractions containing V_1_ were pooled, combined with 500 µl of 1.6 mg/ml H_chim_, concentrated to 2 ml using a 50,000 MWCO VivaSpin concentrator, and further purified over a Superdex200, 1.6 cm x 50 cm size exclusion column. Fractions containing V_1_ were pooled and concentrated and the final concentration was determined using a Nanodrop spectrophotometer set to 280 nm.

### V_o_ expression and purification

V**_o_** was expressed and purified as previously described (Couoh-Cardel et al, 2016; Couoh-Cardel et al, 2015). Briefly, yeast strain YSC1178-7502926 containing a calmodulin-binding protein (CBP) tag at the C-terminus of Vph1p (subunit *a*) was grown in 50 ml high-glucose YPD medium (yeast extract, peptone, 4% glucose) for 15 hours at 200 rpm to inoculate 1 l of high-glucose YPD medium, which was then incubated at 30 °C at 200 rpm for 8 hours. At OD_600_ 12-13, the 1 l culture was split into 12 x 1 l high-glucose YPD medium, bringing the starting OD_600_ to 0.4. The 12x 1 l cultures were incubated at 30 °C for 16 hours at 200 rpm. The cells were harvested at 4,000 rpm for 25 minutes at 4 °C. Pellets were resuspended in 20 mM Tris-HCl, pH 7.4, 500 mM Sorbitol, 2 mM CDTA and stored at -80 °C. Cells were grown in this way until obtaining ∼320 g cells (∼30 l). For preparation of membranes, 320 g of cells were thawed and lysed using 500 ml zirconia beads (Biospec, Inc.) in an inverted bead-beater (chilled in an ice/rock salt bath) for 8 x 1 min at 20,000 rpm separated by 5-6 minute cool-downs at 1,000 rpm. Cell debris and mitochondria were pelleted at 3,000 and 12,500 rpm, 4 °C for 10 min each, respectively, and membranes were collected by centrifugation at 60,000 rpm for 2 hours in a 70Ti ultracentrifuge rotor. Membrane pellets were resuspended in 20 mM Tris-HCl, 500 mM sorbitol, pH 7.4 and stored at -80 °C. The membrane protein concentration was determined using a BCA-TCA assay. For V_o_ purification, two grams of membranes were thawed and brought to 0.6 mg/mg n-undecyl-β-D-maltopyranoside (UnDM). The membranes were gently mixed for 30 min at 4 °C before being brought to 4 mM CaCl_2_. The membranes continued gently mixing for another 30 minutes at 4 °C before being centrifuged at 42,000 rpm for 1 hour. The supernatant was passed over a 15 ml calmodulin column pre-equilibrated with 5 CV calmodulin washing buffer (CWB) containing 10 mM Tris, 100 mM NaCl, 2 mM CaCl_2_, pH 7.4. The column was washed with ∼7 CV CWB, followed by ∼7 CV CWB (sans NaCl), before elution of V_o_ in ∼7 CV 10 mM Tris, 100 mM NaCl, 10 mM CDTA, pH 7.4. Fractions were analyzed by SDS-PAGE, and fractions containing V_o_ were combined and concentrated in a 100,000 MWCO VivaSpin concentrator. The concentration of V_o_ was determined with the BCA-TCA assay.

### V_o_ reconstitution into lipid nanodisc

V_o_ was reconstituted into lipid nanodisc as previously described (Stam & Wilkens, 2016). Briefly, V_o_, MSP, *E. coli* polar lipid extract (Avanti), and UnDM were mixed in this order in the following weight ratios: 1 mg V_o_:2 mg MSP:2.4 mg EPL:1.5% of the reaction volume UnDM. Components were incubated for 1 h on a rotating mixer at room temperature. Detergent was removed over the course of 2 hours on a rotating mixer at room temperature using 0.4 g/ml SM- 2 BioBeads (Bio-Rad). V_o_ND was separated from the beads by centrifugation, and an excess of 2 mM CaCl_2_ over CDTA was added. V_o_ND was captured on a 3 ml calmodulin column and washed with 5 CV CWB before eluting with 10 mM Tris-HCl, 100 mM NaCl, 10 mM CDTA, pH 7.4. in 6 x 1 ml fractions. V_o_ND containing fractions were combined and concentrated to 2.2 ml using a 100,000 MWCO VivaSpin concentrator. The concentration of V_o_ND was determined using a Nanodrop spectrophotometer set to 280 nm.

### Reconstitution of V_1_(H_chim_)V_o_ND

Functional reconstitution of V_1_H_chim_ and V_o_ND was achieved as described (Khan et al, 2022; Sharma et al, 2019). V_1_H_chim_, V_o_ND, and C were combined in a 1:1:3 molar ratio, respectively. Samples were incubated at room temperature for 2 hours or overnight at 18 °C, and then stored at 4 °C.

### Nanobody expression and purification

Nanobodies (Nbs) containing C-terminal 7x His and HA tags were received from the VIB Nanobody facility in *E. coli* TG1 with pMECs-GG vector containing nanobody genes. Nbs were transformed into WK6 *E. coli* for expression (pMECs vector with C terminal HA-His tags). A 5 ml RB (LB + 0.2% glucose) starter culture supplemented with 50 µg/ml Carbenicillin was inoculated with Nb expressing *E. coli* and grown at 37 °C and 180 rpm overnight. 1 l Terrific Broth (LB + glucose + glycerol) + 1 mM Carbenicillin was inoculated with 3 ml of the overnight starter culture. The cells were grown to OD_600_ 0.6-0.9 before induction with 1 mM IPTG. The cells were grown for 15 hours at 28 °C at 200 rpm post induction before harvesting at 3,000 rpm, 4 °C for 25 min. Pellets were resuspended in 20 ml buffer A (20 mM Tris-HCl, pH 8, 250 mM NaCl, 10 mM imidazole), and stored at -20 °C until use. Cell pellets were thawed and lysed by sonication 3x for 30 seconds with 30 second breaks in-between on ice. Cell debris was removed using centrifugation at 13,000 x g, 4°C and supernatant applied to a Ni-NTA column, washed, and eluted using a linear imidazole gradient (10 - 500 mM). Nb containing fractions were pooled and concentrated using a 5000 MWCO VivaSpin concentrator and applied to a Sephadex-75 1.6 cm x 50 cm gel filtration column attached to an ÄKTA FPLC. Nbs were concentrated and stored at 4 °C or -80 °C for long term storage. Final protein concentration was determined using UV-absorbance at 280 nm.

### High-throughput expression and purification of nanobodies

Expression level of Nbs in *E. coli* WK6 was tested by growth and purification using 48- well plates. Briefly, 5 ml Rich Broth supplemented with 5 µl 50 mg/ ml carbenicillin and 5 µl nanobody stocks were grown overnight at 37 °C at 180 rpm. The next day, Nbs were grown in 750 µl Terrific Broth supplemented with 50 µg/ml carbenicillin and 1mM IPTG per well in a 48- well plate was inoculated with 5 µl of the overnight Nb cultures. The 48-well plate was grown in a TECAN for 18 hours at 28 °C with shaking. The plate was centrifuged at 4,000 rpm for 10 minutes at 4 °C to pellet cells. Media was removed by aspiration and plates were stored at -80 °C until use. Plates were thawed at room temperature, andwhile100 µl Bacterial Protein Extraction Reagent (BPER, Pierce) was warmed to room temperature. BPER was brought to supplemented with 1 mM PMSF added per well and cells were incubated with 100 µl per well of BPER for 20 minutes at room temperature at with shaking at 300 rpm. 400 µl of 1x PBS (400 µl) was then added to each well along with 7.5 µl of magnetic Ni-NTA beads (GoldBio), and incubated at room temperature for 1.5 hours at 300 rpm. For all washes and elution, magnetic beads were immobilized using a home-made magnetic board to remove the solution via aspiration. Lysate was removed by aspiration and 500 µl 1x PBS was added to each well and incubated, not letting the wells dry out. The plate was returned to the shaker (300 rpm) for 20 minutes at room temperature, and this washing process was repeated thrice. After the third PBS wash, 100 µl 35 mM imidazole (in 1x PBS) was added to each well and incubated with the beads for 10 minutes at room temperature at 300 rpm. After 10 minutes the magnetic board was added to the shaker under the 48-well plate, and continued shaking at 300 rpm at room temperature for 10 minutes. At this point the magnetic board remained under the 48-well plate for the rest of the protocol. 500 µl 1x PBS was added to each well and the plate continued shaking at 300 rpm at room temperature for 15 minutes. The liquid in each well was removed via aspiration and the Nbs were eluted off the beads using 65 µl of 250 mM Imidazole in 1x PBS on the shaker at 300 rpm at room temperature for 20 minutes. Nbs were run on 15% SDS-PAGE gels to verify expression levels, and stored at 4 °C. Expression levels were quantified using FIJI (**Appendix Fig. S1**).

### ATPase Assays

ATPase assays were done with an ATP regenerating coupled enzyme system containing 5 mM ATP, 0.5 mM NADH, 2 mM phosphoenolpyruvate, and 30 units/ml each of pyruvate kinase and lactate dehydrogenase, 50 mM HEPES pH 7.5, at 37°C as described in (Oot, 2016). For determining Nbs’ impact on V-ATPase activity, Nbs were added to 10 µg *in vitro* reconstituted V_1_(H_chim_)V_o_ND and incubated for at least 30 min at 20-25 °C before measuring the activity. For measuring assembly inhibition, V_o_ND was pre-incubated with Nb and subunit C for 30 min before adding V_1_(H_chim_).

### Biolayer Interferometry (BLI)

A 16 channel BLI instrument (ForteBio Octet Red384) was used to determine binding affinity of Nbs for V_o_ND. Ni-NTA sensors were hydrated in 20 mM Tris-HCl, 100 mM NaCl, 5 mg/ml BSA, pH 7.1-8 (BLI buffer) for 10 min prior to collecting baselines for 300 seconds in the same buffer. Ni-NTA sensors were loaded with 0.02 mg/ml Nb followed by a baseline of 300 seconds in BLI buffer and association with V_o_ND for 2400 seconds. A titration of 60 nM to 0.75-0.083 nM V_o_ND (depending on experiment) in BLI buffer was used. Dissociation in BLI buffer was monitored for 3600 seconds. To correct for non-specific binding of V_o_ND to empty Ni-NTA sensors, non-specific binding reference controls for each V_o_ND concentration were run in parallel and subtracted from sample sensors pairwise. Data analysis was performed in GraphPad Prism 9 using a Robust Fit of Association and Dissociation data that had been corrected for non-specific binding.

### Cryo electron microscopy

For cryo electron microscopy (cryoEM) structure determination, Nb bound V_1_(H_chim_)V_o_ND or V_o_ND at 1-3 mg/ml was vitrified on freshly glow discharged C-Flat or Au-Flat 1.2/1.3 grids (Protochips) in liquid ethane using a self-built plunger operating in a 4 °C cold room, or a Leica EM GP2 set to 6°C and 90% humidity operating in sensor blot mode. Grids were examined using a ThermoFisher Glacios transmission electron microscopes equipped with Falcon IVi direct detector. Data for N27 bound V_1_(H_chim_)V_o_ND were recorded at the cryoEM facility at the Department of Biological Sciences, Seoul National University, Seoul, Korea. Data for N125 and N2149 were obtained from the cryoEM facility at the Hauptman-Woodward Institute in Buffalo, NY, USA. For N27, a dataset of 3,215 movies was initially analyzed using the cryoSPARC package of programs (Punjani et al, 2017) (**Appendix Figure S4**; **Appendix Table S1**). To resolve heterogeneity of rotary states, the dataset was analyzed using the RELION 5 package of programs (Zivanov et al, 2018) (**Appendix Figures S5,6**; **Appendix Tables S2,3**).

### Model building

Model building for Nb bound V_o_ND complexes was started using our previous 2.7 Å structure of V_o_ND (PDBID: 6m0r; (Roh et al, 2020)). Coordinate files for the Nbs were obtained from the Phyre2 (Kelley et al, 2015) or Alphafold (Jumper et al, 2021) servers. Atomic models were placed into the cryoEM density using rigid body fitting in Chimera (Pettersen et al, 2004) followed by manual building and automated real space refinement using Coot (Emsley et al, 2010), ISOLDE (Croll, 2018), and Phenix (Adams et al, 2010). Models for N27 bound state 1, 2’ and 3’ V_1_(H_chim_)V_o_ND were obtained by rigid body fitting of subcomplexes and subunits from models of state 1 and state 3 V_1_(H_chim_)V_o_ND (PDBID: 7fda,c, (Khan et al, 2022)) and the model of the N27 bound V_o_ND reported here (PDBID: 9e7l).

### Generation of BAP- (Avi)-tagged nanobodies

Nanobody in pMECs vector was used as a template for PCR amplification and restriction-based cloning (NotI and PstI) into a modified (Genscript) pET28a vector to contain a C-terminal 7x His tag, Avi tag, and PPase cleavage site (HBP), with the cleavage site being closest to the C-terminus of the Nb. The primers used are as follows: Nb_PstI_intoHBPF: CAGGTGCAGCTGCAGGAGTCTG and Nb_NotI_intoHBPR: TTACCATGCGGCCGCTGAGGAGA. N125^HBP^ in the modified pET28a vector, with kanamycin resistatnce, was co-transformed with a vector containing BirA, with chloramphenicol resistatnce, into BL21DE3 cells. Cells were grown in 20 ml Rich Broth (LB+ glucose), with 1 mM kanamycin and chloramphenicol overnight at 37 °C. The overnight culture was transferred to 1 l LB supplemented with 100 µM Biotin, 100 µM NaH_2_PO_4_, 1 mM kanamycin, and 1 mM chloramphenicol and grown at 37 °C and 200 rpm until the OD_600_ was in between 0.6 and 0.9 (∼ 1 h). Cells were induced with 1 mM IPTG, and grown overnight at 28 °C, 200 rpm. The next morning cells were pelleted at 3000 rpm, 4 °C, for 25 min. Pellets were resuspended in 20 ml Nb Buffer A (see the above protocol for purification of Nbs), and stored at -80 °C until use. Nbs were purified as previously outlined.

### NeutrAvidin-binding to N125^HBP^

NeutrAvidin (ThermoFisher) was dissolved into 1 mg/ml working stock in water. NeutrAvidin was incubated with a four-fold molar excess of N125^HBP^ for 1 h at room temperature. The mixture was then incubated on 15 µl streptavidin beads for 1 h at room temperature on a rotor to remove any unbound Nb. After the incubation, the beads were spun down at 1 rcf, for 2 min, at room temperature and the supernatant was removed and set aside for use.

### Purification of *Hs*V_1_V_o_

Human V-ATPase (*Hs*V_1_V_o_) was purified as previously outlined in (Oot et al, 2021). Briefly, *HsV1Vo was purified from HEK 293F cells stably expressing* C-terminally 2xFLAG tagged *a* subunit, isoform 4, or *a*4, Cells were maintained at 37 °C in a humidified, 8% CO2 atmosphere in Gibco Freestyle media. Cells were collected in 1.6 l batches by centrifugation at 300 x g, 15 min, 4 °C and pellets flash frozen on liquid nitrogen and placed at -80 °C until use. For purification, pellets were thawed at room temperature and lysed via dounce homogenization, followed by centrifugation at 720 x g for 5 minutes at 4 °C. The supernatant was set aside, while the pellet was resuspended in TBSE (20 mM Tris-HCl, 150 mM NaCl, 0.5 mM EDTA, pH 7.2) containing protease inhibitors (4 mg/ml leupeptin, 4 mg/ml pepstatin, 1 mg/ml chymostatin and 1mM PMSF), and processed by another round of dounce homogenization and centrifugation as above. The supernatants were combined and centrifuged at 14,629 x g at 4 °C for 1 h, and the pellet was resuspended in 8 ml of 15 mg/ml MSP containing protease inhibitors (4 mg/ml leupeptin, 4 mg/ml pepstatin, 1 mg/ml chymostatin and 1mM PMSF). Dodecyl maltopyranoside was added to 1%, and the sample incubated with rotation, 45 min at 4 °C after which 0.4 g of BioBeads per ml of material was added to the solubilized pellet to remove detergent, and the mixture continued to rotate at 4 °C for another 2 h. The sample was removed from BioBeads by centrifugation, and added to 1.5 ml FLAG beads pre-equilibrated with TBSE and protease inhibitors for batch binding 1.5 h, 4 °C before applying the sample and beads to a column for subsequent washing and elution. The column was washed with 50 ml TBSE containing protease inhibitors, and eluted in TBSE containing 250 μg / ml 3x FLAG peptide and protease inhibitors, in 1 CV fractions. *Hs*V_1_V_o_ ATPase activity was measured as in (Oot et al, 2021).

### Pulldown of *Hs*V_1_V_o_

30 µl of a 50% suspension of streptavidin beads (15 µl settled beads) were aliquoted into 3 separate tubes. Beads were equilibrated by washing three times in 45 µl TBSE (20 mM Tris-HCl, 150 mM NaCl, 0.5 mM EDTA, pH 7.2). Biotinylated, BAP-tagged N125 (20 µg; N125^HBP^) was loaded onto the beads in 2 of the 3 tubes and incubated for 1 hour at room temperature while rotating, with the third tube receiving only TBSE instead of Nb. The beads were pelleted by centrifugation at 1000 x g, for 2 min, at room temperature and supernatant collected (FT). The beads were then washed 3 times with 45 µl TBSE, pelleted as above, before 10 µg of *Hs*V_1_V_o_ was mixed into the beads of 2 of the 3 tubes, one containing streptavidin bound N125^HBP^ and the other containing just beads. The tube containing just streptavidin bound Nb received TBSE instead. The beads were incubated for 1 hour at room temperature while mixing, and then pelleted at the standard conditions. The supernatant from each tube was saved (FT). Each tube was then eluted in 30 µl 1xTBSE with 2 mM DTT and 2 µl 1.68 mg/ ml PPase at 4 °C overnight while rotating. The beads were then pelleted at the standard conditions and the eluate was saved from each tube. 12 µl of each saved sample was resolved on 15% SDS-PAGE gels, with 3 µl of sample loading buffer containing 60 mM DTT. The gels were then silver stained to visualize the protein.

## Acknowledgements

This research was supported by NIH grant GM141908 to S.W. and grants 2020R1A6C101A183 and 2021M3A9I4021220 from the National Research Foundation, Republic of Korea to S.-H.R. We thank the Director and staff of the VIB Nanobody core at the Vrije Universiteit Brussel, Brussels, Belgium for generating the anti-yeast V_o_ND Nbs and for helpful discussions, and the staff of the Hauptman-Woodward Institute cryoEM facility for cryoEM data collection.

## Author contributions

K.K. conducted biochemical and biophysical characterization of Nb-V-ATPase interactions and assisted with cryoEM structure determination. J.B.P. collected cryoEM data for N27 bound to V_1_(H_chim_)V_o_ND and performed image reconstruction in cryoSPARC supervised by S.-H.R. R.A.O. provided human 293 suspension cells and purified human V-ATPase, assisted by K.K. Md M.K. purified yeast V_o_ND for Nanobody generation. S.W. supervised the project and conducted cryoEM structure determination. K.K. wrote the first draft of the manuscript with input from S.W. and editing by R.A.O, Md M.K., and J.B.P.

## Data availability

Atomic coordinates and maps for N27:V_o_ND, N125:V_o_ND and N2149:V_o_ND have been deposited under 9e7l, 9e76, 9mj4, and EMDB-4767, EMDB-47659, and EMDB- 48311, respectively. Maps for V_1_(H_chim_)V_o_ND:N27 and focused refinement of the V_o_ND regions for state 1, state 3^+18^ and state 3^-18^ are deposited under EMD-48593, EMDB-48594, EMDB- 48592, EMDB-48595, EMDB-48597, and EMDB-48596, respectively.

## Appendix Figures

**Appendix Figure S1:**
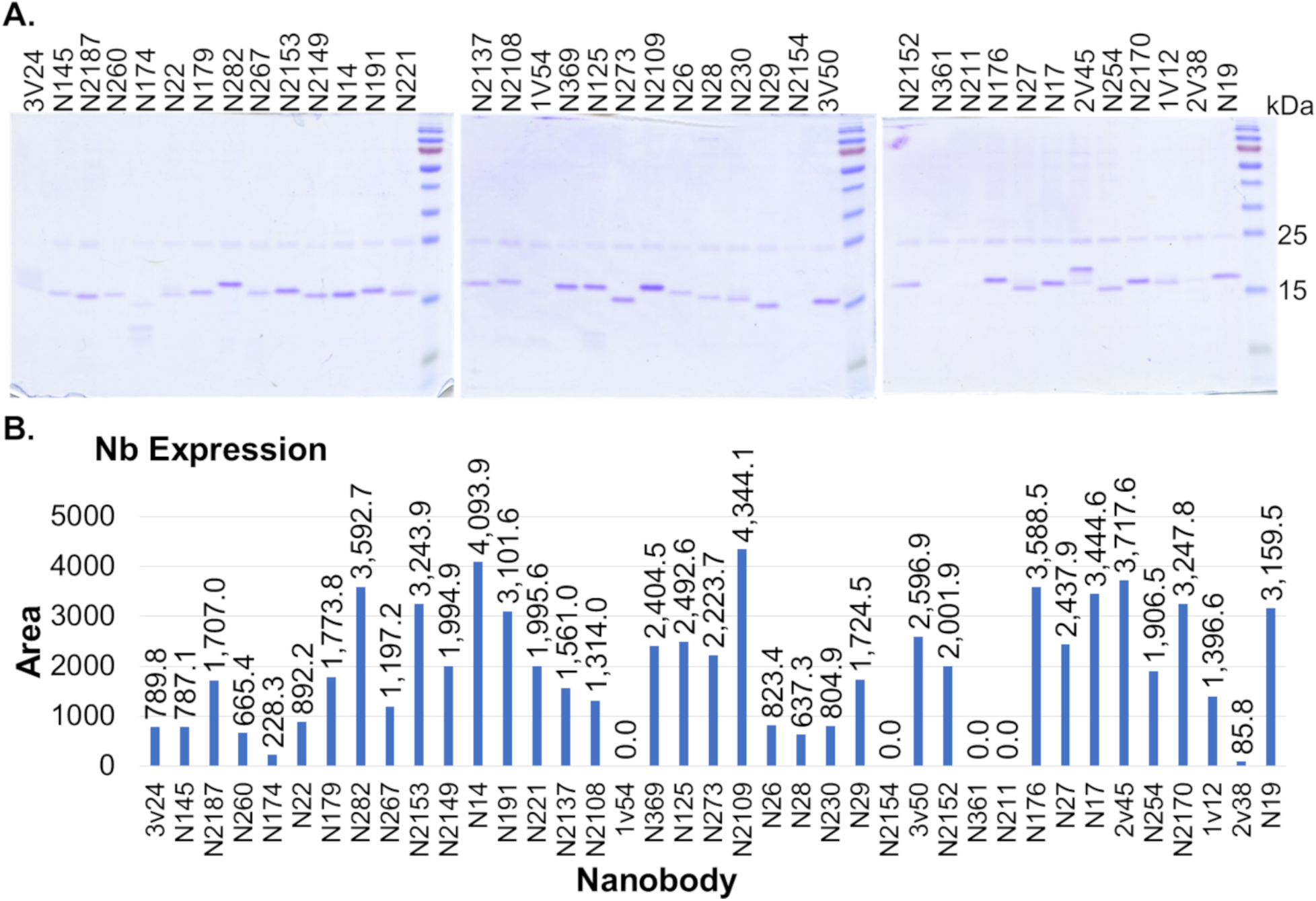
High-Throughput Nanobody Expression. **A** Each purified nanobody was resolved on a 15% acrylamide gel to verify expression. **B** Expression was quantified on FIJI by graphing a vertical cross-section of band intensity for each lane containing a nanobody, and calculating the area under the curve.

**Appendix Figure S2:**
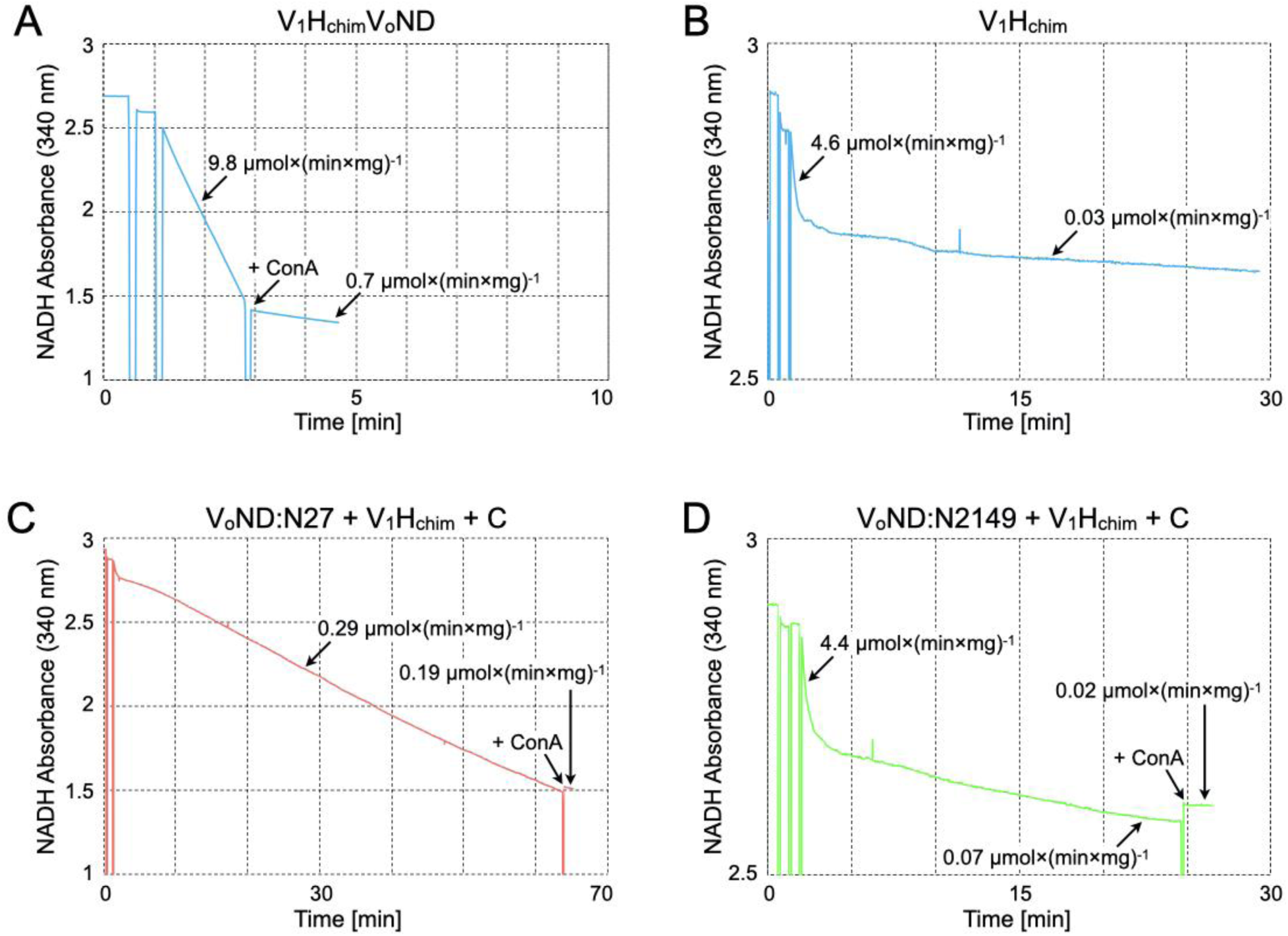
V-ATPase activity traces. **A** Representative ATPase activity trace for *in vitro* reconstituted V_1_(H_chim_)V_o_ND. At the time indicated by the arrow, 200 nM of the V-ATPase specific inhibitor Concanamycin A (ConA) is added. **B** Representative ATPase activity trace for V_1_H_chim_. The activity decays quickly due to trapping of inhibitory MgATP in a catalytic site. **C** Assembly of N27 bound V_o_ND with V_1_H_chim_ and subunit C. Note the different time scale compared to panel (A). **D** Assembly of N2149 bound V_o_ND with V_1_H_chim_ and subunit C. Note that the activity profile resembles that of isolated V_1_H_chim_, indicating that N2149 binding prevents holoenzyme assembly.

**Appendix Figure S3:**
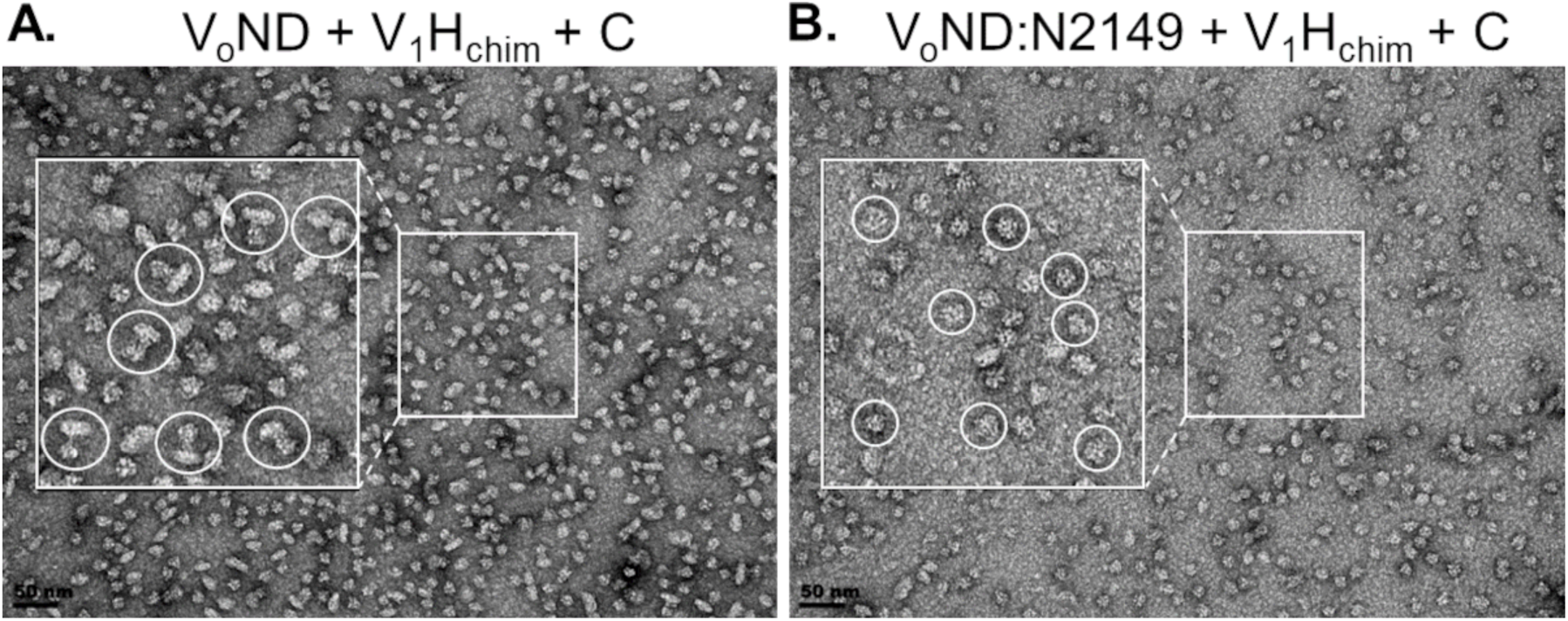
Negative stain EM of holoenzyme assembly mix in presence and absence of N2149. **A** *In vitro* assembly of V_o_ND with V_1_H_chim_ and subunit C produces a significant population of holo V_1_(H_chim_)V_o_ND characterized by their dumbbell shape as highlighted by the white circles. Bar is 50 nm. **B** Upon reconstitution in the presence of N2149, only individual V_1_H_c_ and V_o_ND particles are observed, characterized by their globular shape and highlighted by the white circles. Bar is 50 nm.

**Appendix Figure S4:**
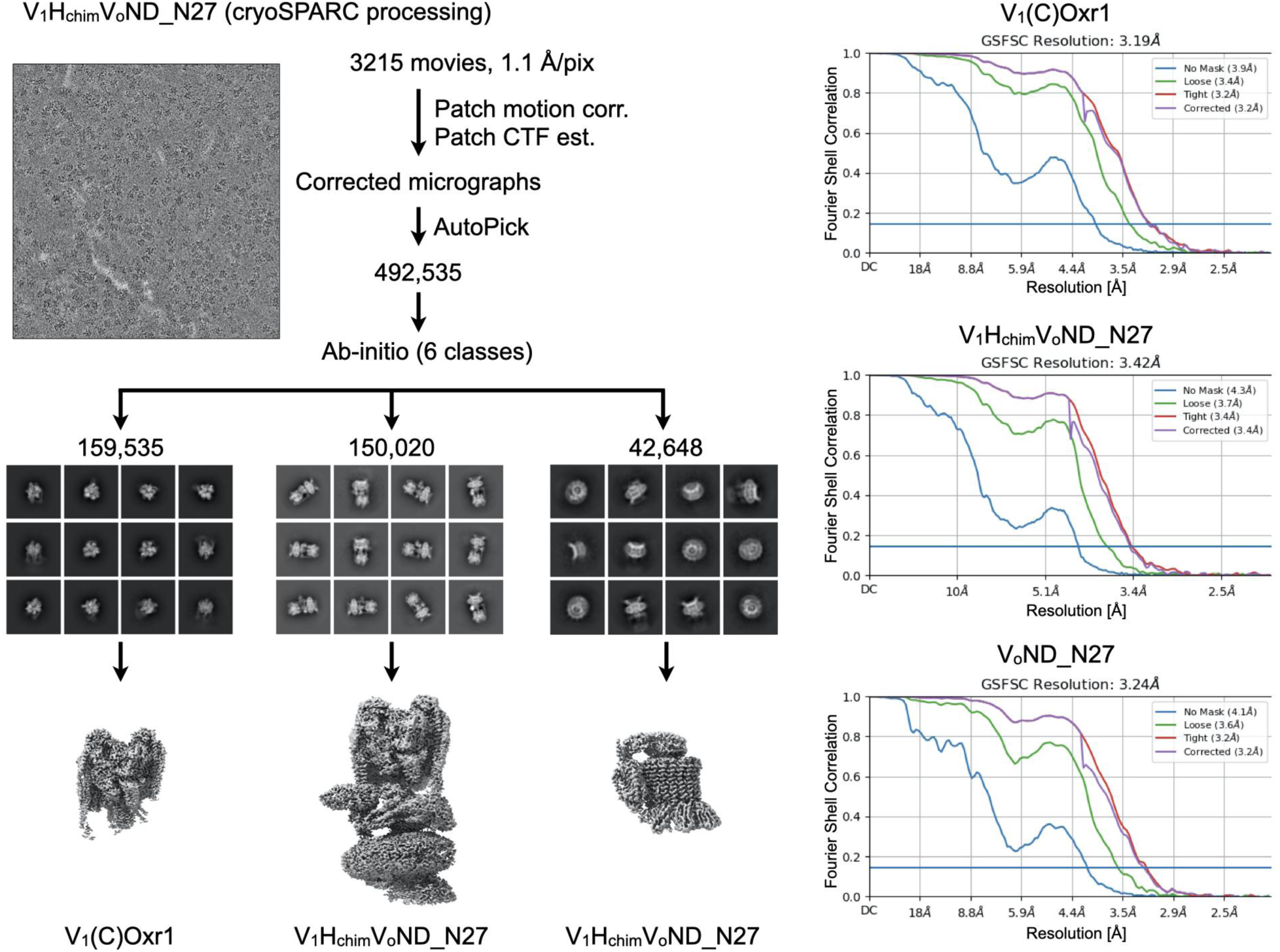
**Cryo-EM analysis of N27 bound V_1_(H_chim_)V_o_ND in cryoSPARC**

**Appendix Figure S5:**
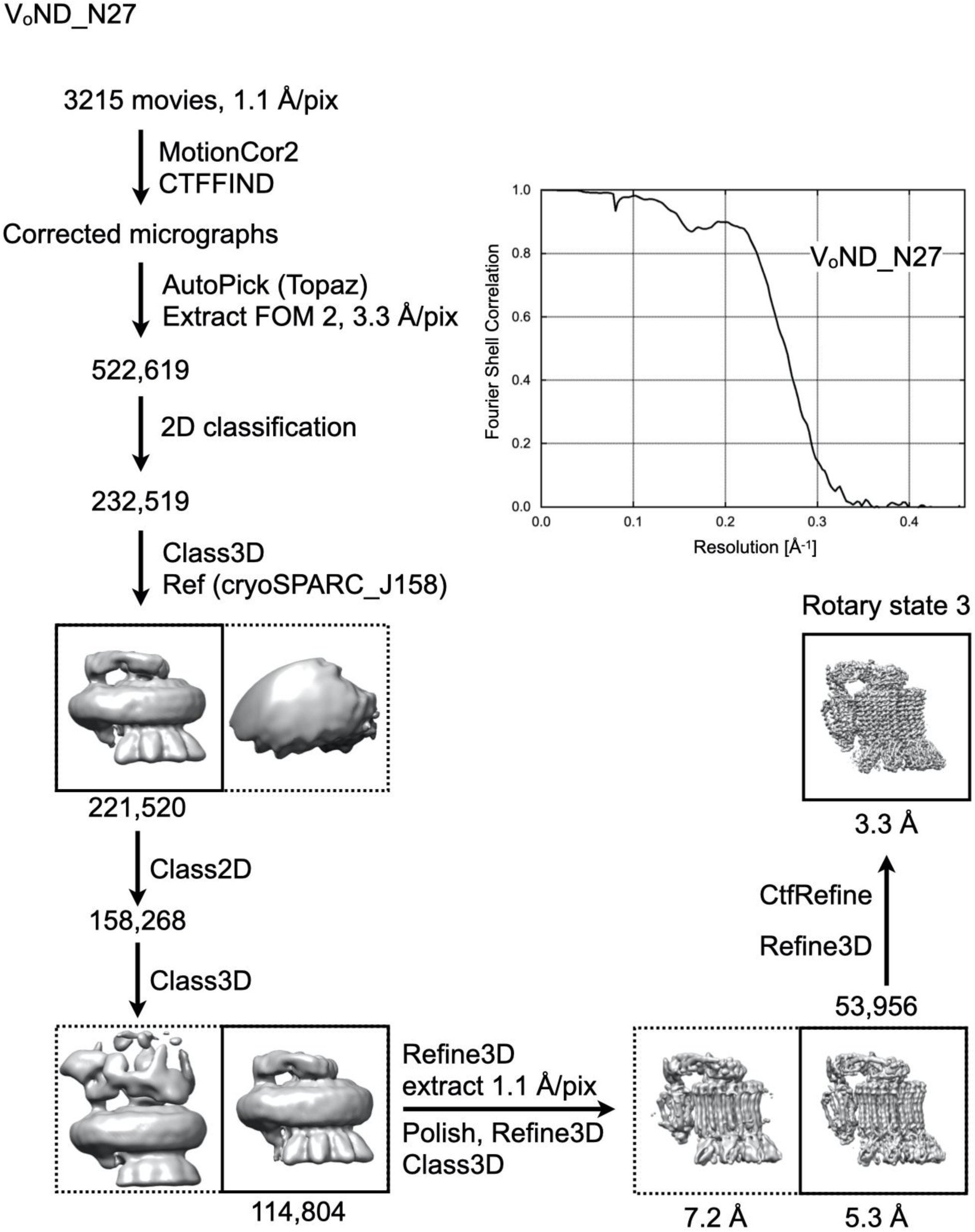
**Cryo-EM analysis of N27 bound V_o_ND in RELION 5**

**Appendix Figure S6:**
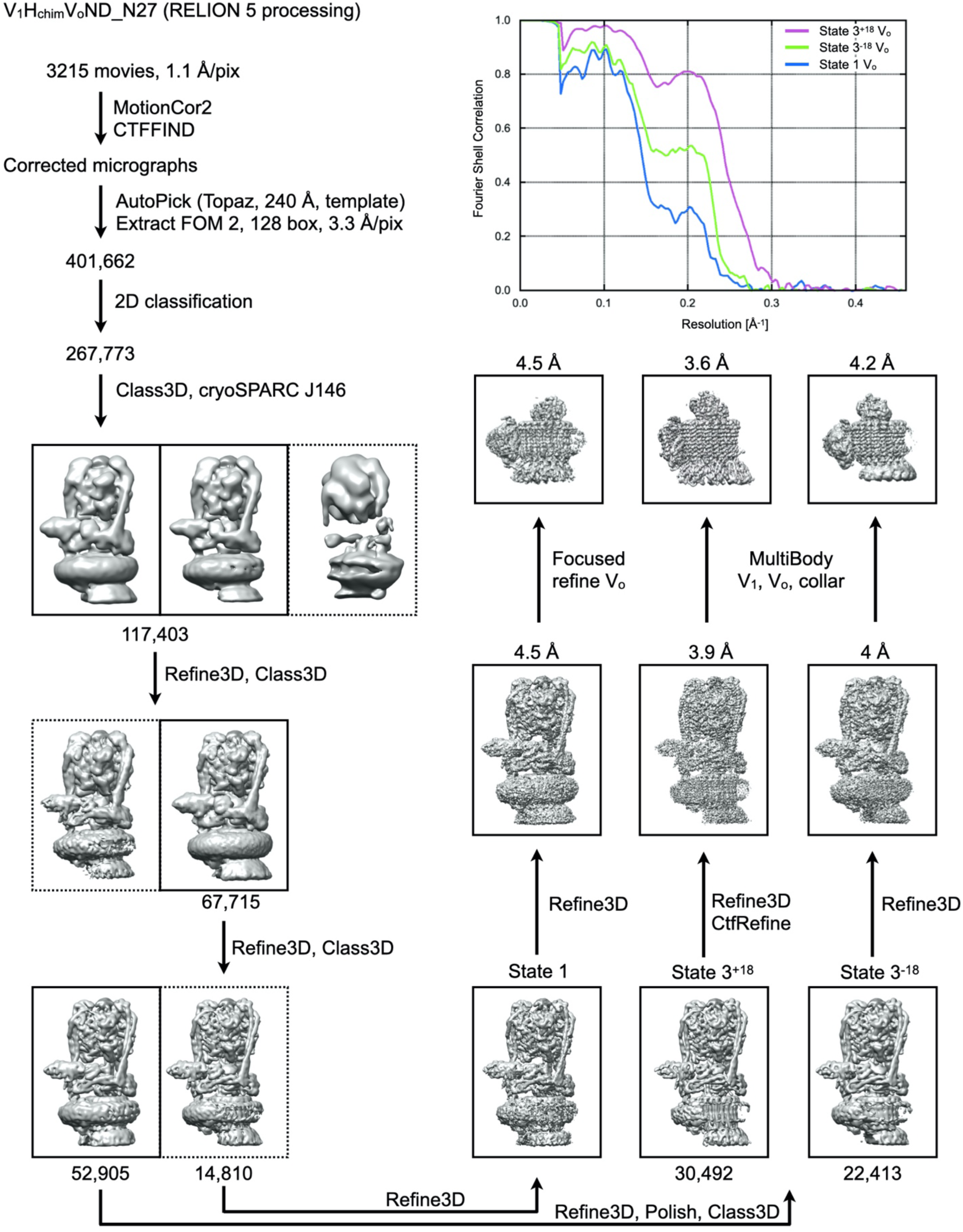
**Cryo-EM analysis of N27 bound V_1_(H_chim_)VoND in RELION 5**

**Appendix Figure S7:**
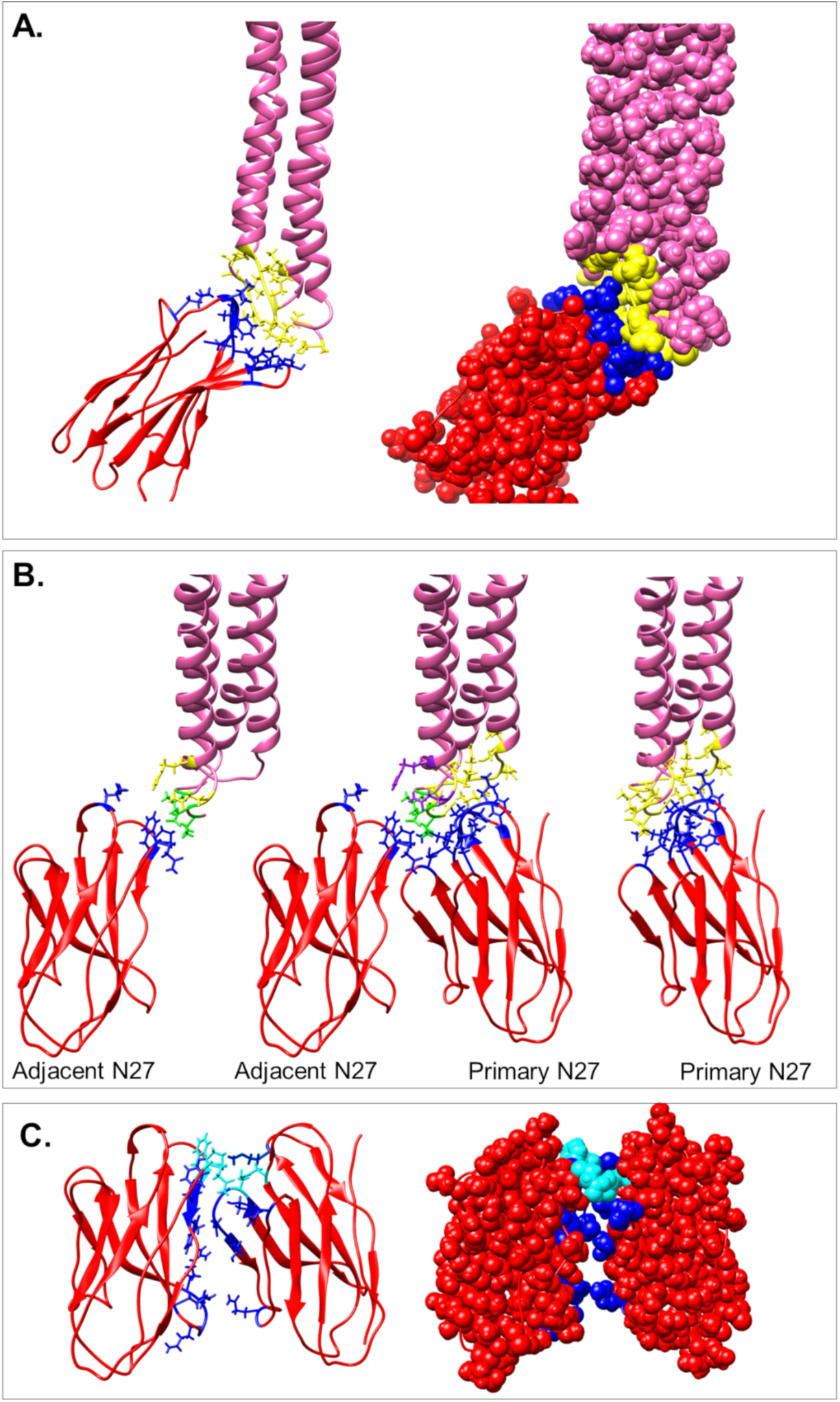
N27:*c* subunit interfaces as determined by PISA. **A** Amino acids of N27 (red) that contact residues in *c* are shown in blue. Amino acid residues of *c* (pink) that are in contact with N27 are shown in yellow. **B** Amino acids of *c* (pink) that contact the primary N27 are shown in yellow, and contacts with the adjacent N27 are shown in purple. Residues shown in green were buried in both the primary N27 and adjacent N27 separately. **C** N27 to N27 contacts are highlighted in dark blue, with residues contacting both *c* and a neighboring Nb in cyan.

**Appendix Figure S8:**
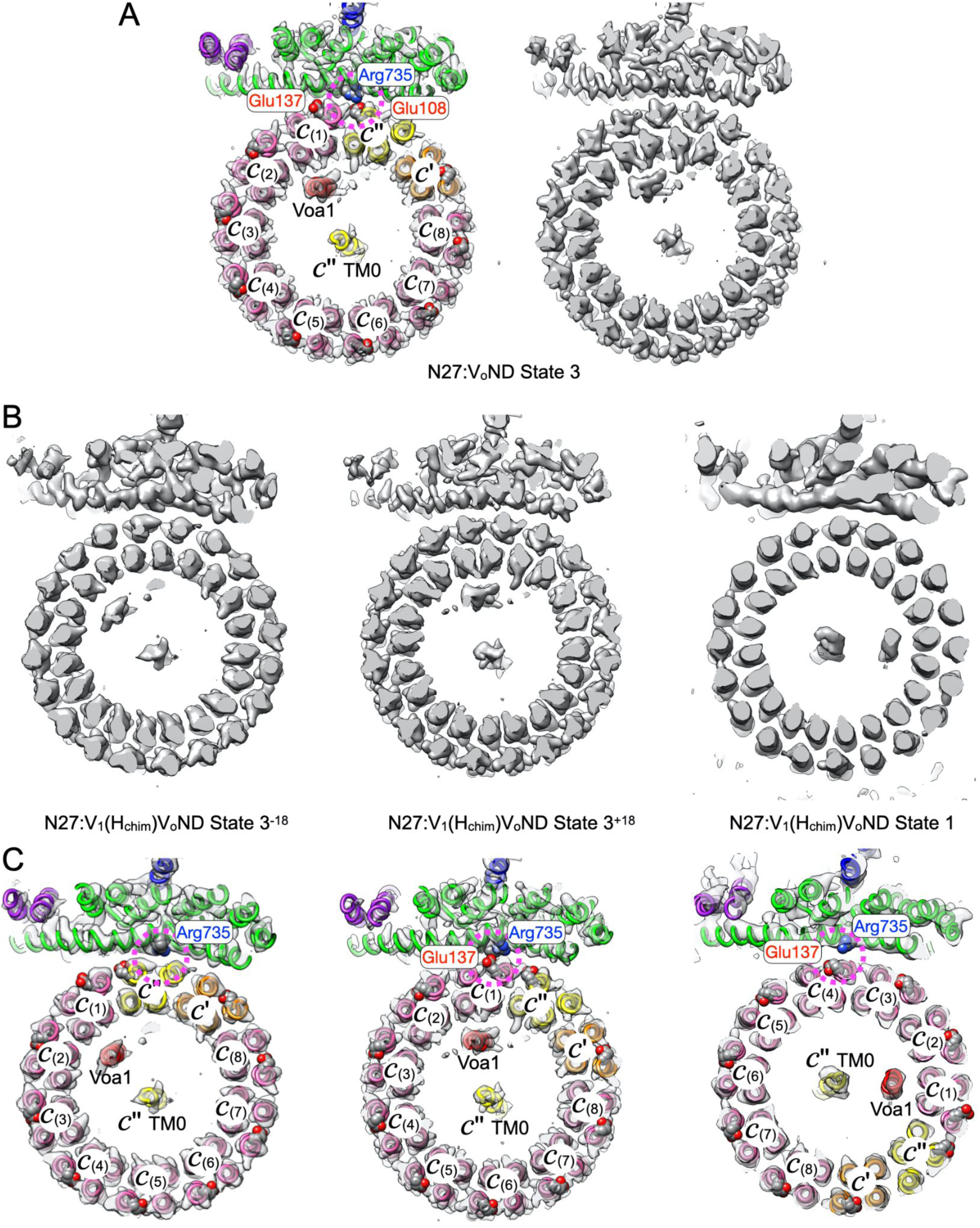
Focused refinement of V_o_ subcomplexes of N27-bound holoenzymes in rotary states 3^-18^, 3^+18^ and 1, and comparison to N27 bound V_o_ND in state 3. **A** Left panel: Cross section of V_o_ND bound to N27 at the level of the conserved *c*-ring glutamates. In the canonical state 3, conserved Arg735 of *a*_CT_ is in contact with Glu108 of *c*’’ (see pink dashed circle). Right panel: EM density. **B** EM density of cross section as in (A) for N27 bound V_1_(H_chim_)V_o_ND in state 3^-18^, state 3^+18^, and state 1. **C** Cross sections as in (B) with rigid body fitted coordinates from 7FDC (Khan et al, 2022) for states 3^-18^ and 3^+18^, and 7FDA for state 1.

**Appendix Figure S9:**
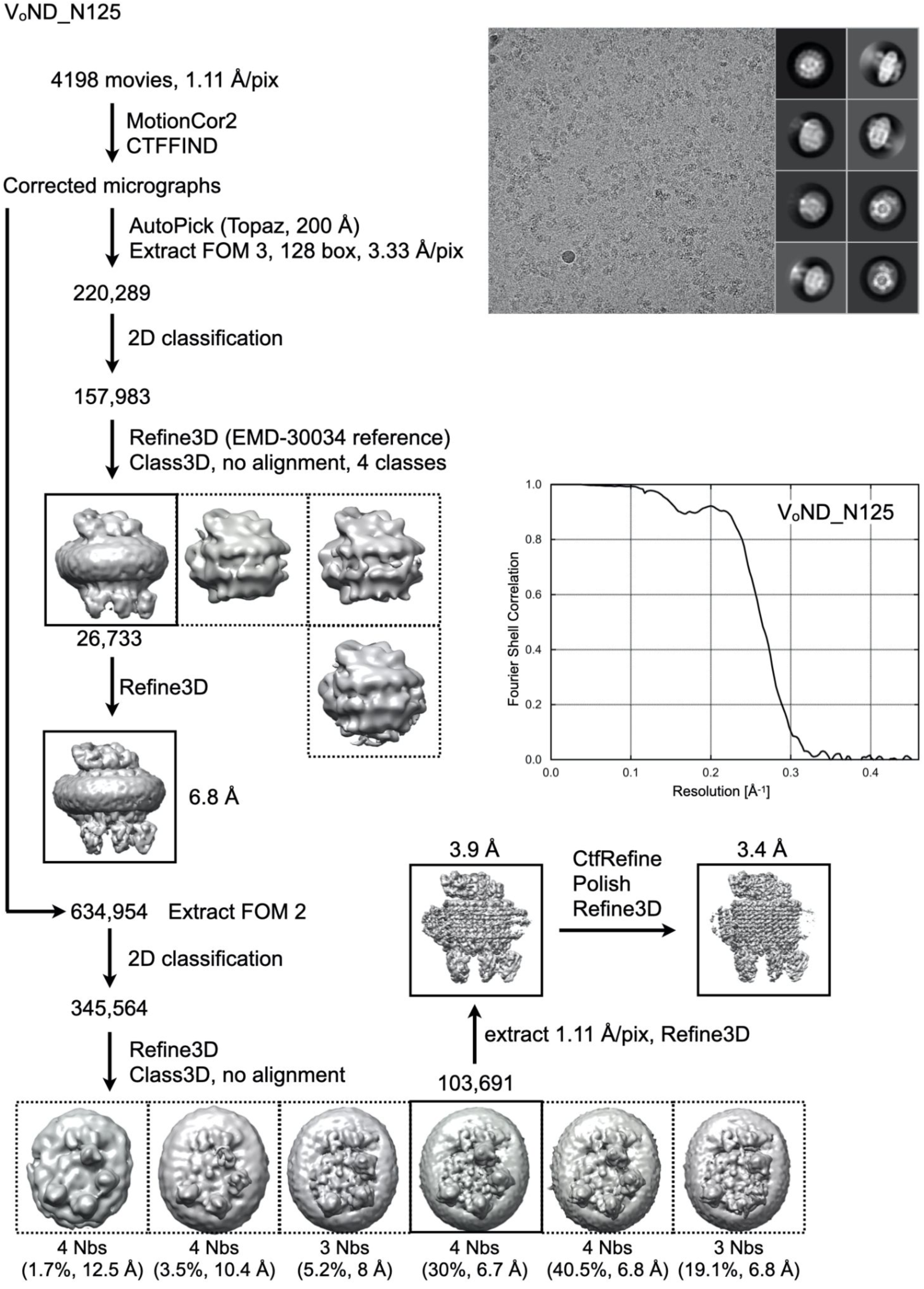
**Cryo-EM analysis of N125 bound VoND**

**Appendix Figure S10:**
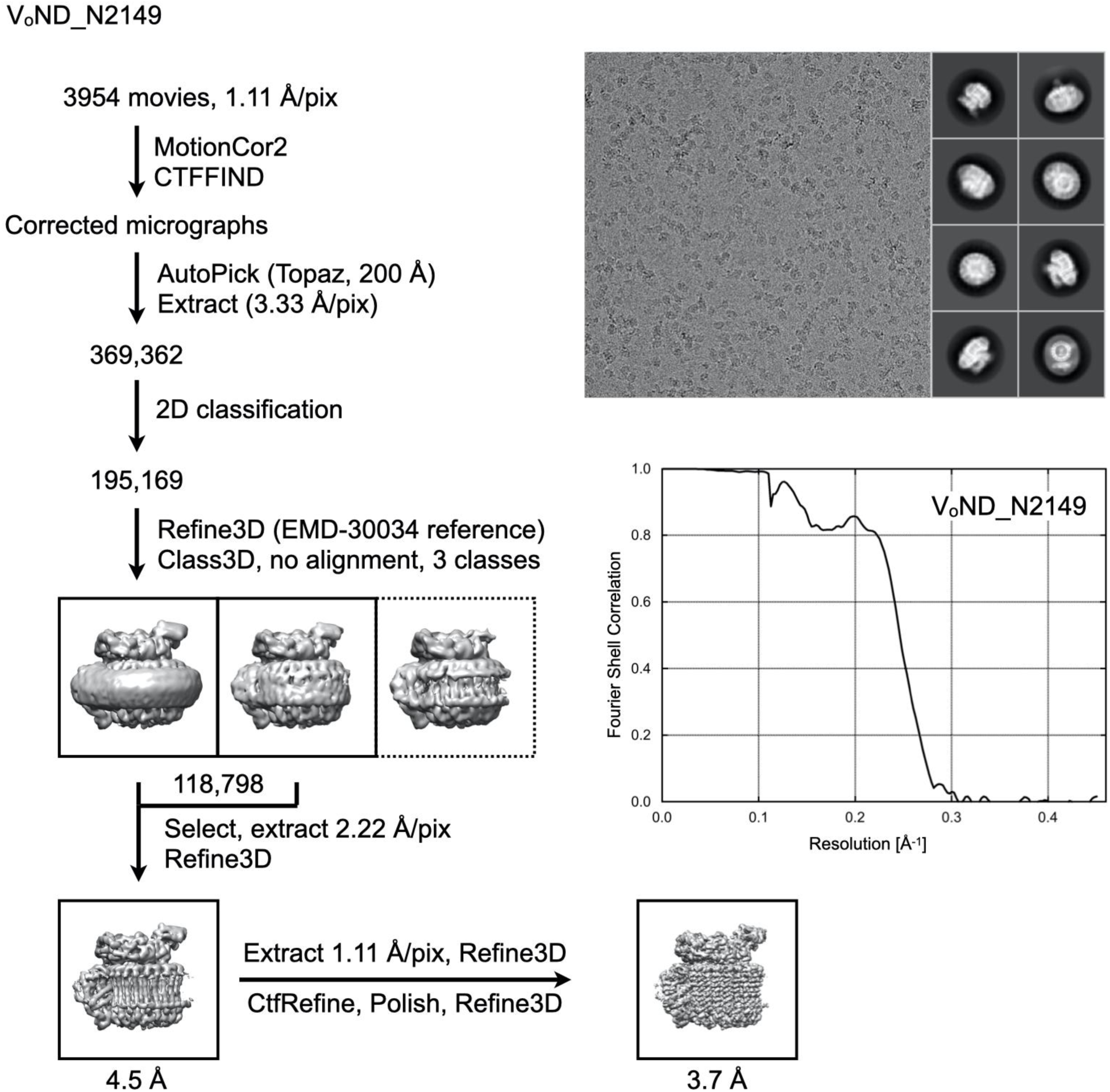
**Cryo-EM analysis of N2149 bound VoND**

**Appendix Figure S11:**
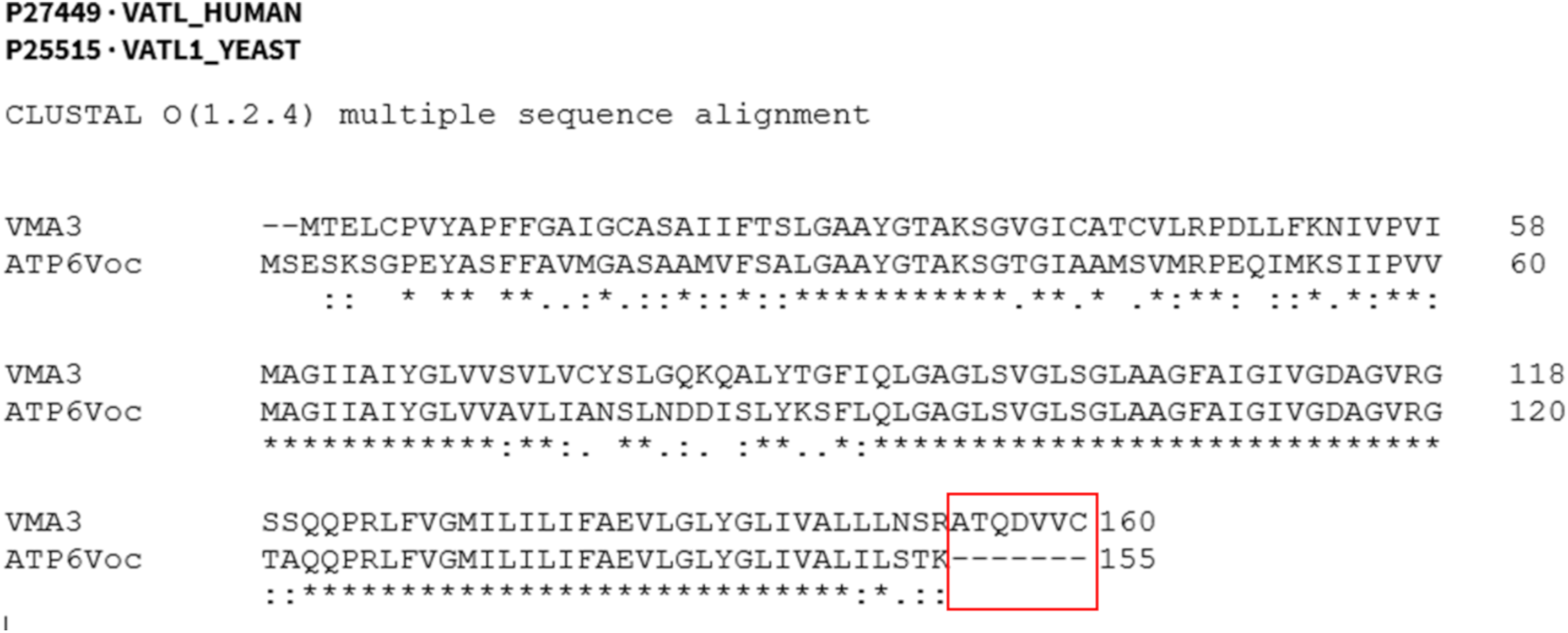
Sequence comparison of human (ATP6V0c) and yeast (VMA3) *c* subunits. Outlined in red are C-terminal residues that are present in yeast, but missing in the human subunit.

**Appendix Table S1.** CryoEM data collection, refinement and validation statistics.

|  | V<sub>o</sub>ND:N27<br>(EMDB-47679)<br>(PDB 9e7l) | V<sub>o</sub>ND:N125<br>(EMDB-47659)<br>(PDB 9e76) | V<sub>o</sub>ND:N2149<br>(EMDB-48311)<br>(PDB 9mj4) |
| --- | --- | --- | --- |
| **Data collection and processing** |  |  |  |
| Magnification | 92,000 | 92,000 | 92,000 |
| Voltage (kV) | 200 | 200 | 200 |
| Electron exposure (e<sup>-</sup>/Å<sup>2</sup>) | 39.96 | 52.5 | 50.3 |
| Defocus range (μm) | 1.3 - 2 | 0.4 - 2 | 0.4 - 2 |
| Pixel size (Å) | 1.1 | 1.11 | 1.11 |
| Movies recorded | 3215 | 4198 | 3954 |
| Symmetry imposed | C1 | C1 | C1 |
| Initial particle images (no.) | 522,619 | 634,954 | 369,362 |
| Final particle images (no.) | 53,956 | 103,691 | 118,798 |
| Map resolution (Å) | 3.33 | 3.4 | 3.7 |
| FSC threshold | 0.143 | 0.143 | 0.143 |
| Map resolution range (Å) |  |  |  |
| **Refinement** |  |  |  |
| Initial model used (PDB code) | 6m0r | 6m0r | 6m0r |
| Model resolution (Å) | 3.4 | 3.4 | 3.8 |
| FSC threshold | 0.5 | 0.5 | 0.5 |
| Model resolution range (Å) |  |  |  |
| Map sharpening *B* factor (Å<sup>2</sup>) | n/a | n/a | n/a |
| Model composition |  |  |  |
| Non-hydrogen atoms | 29,190 | 26,025 | 23,031 |
| Protein residues | 3,839 | 3,423 | 3,035 |
| Ligands | 0 | 0 | 0 |
| *B* factors (Å<sup>2</sup>) |  |  |  |
| Protein (min/max/average) | 52.5/193.2/104 | 54/222.1/104 | 78/169.2/110.9 |
| Ligand |  |  |  |
| R.m.s. deviations |  |  |  |
| Bond lengths (Å) | 0.002 | 0.002 | 0.003 |
| Bond angles (°) | 0.497 | 0.467 | 0.501 |
| Validation |  |  |  |
| MolProbity score | 0.74 | 0.75 | 1.02 |
| Clashscore | 0.75 | 0.78 | 2.4 |
| Poor rotamers (%) | 0.03 | 0.11 | 0 |
| Ramachandran plot |  |  |  |
| Favored (%) | 99.02 | 98.82 | 99 |
| Allowed (%) | 0.98 | 1.09 | 1 |
| Disallowed (%) | 0 | 0.09 | 0 |
| Ramachandran Z-score (RMSD) |  |  |  |
| Whole | 0.87 (0.13) | 1.25 (0.14) | 0.74 (0.15) |
| Helix | 0.92 (0.11) | 1.31 (0.11) | 0.8 (0.11) |
| Sheet | -0.07 (0.21) | -0.81 (0.32) | -0.6 (0.56) |
| Loop | 0.56 (0.19) | 0.42 (0.2) | 0.1 (0.21) |

**Appendix Table S2.**
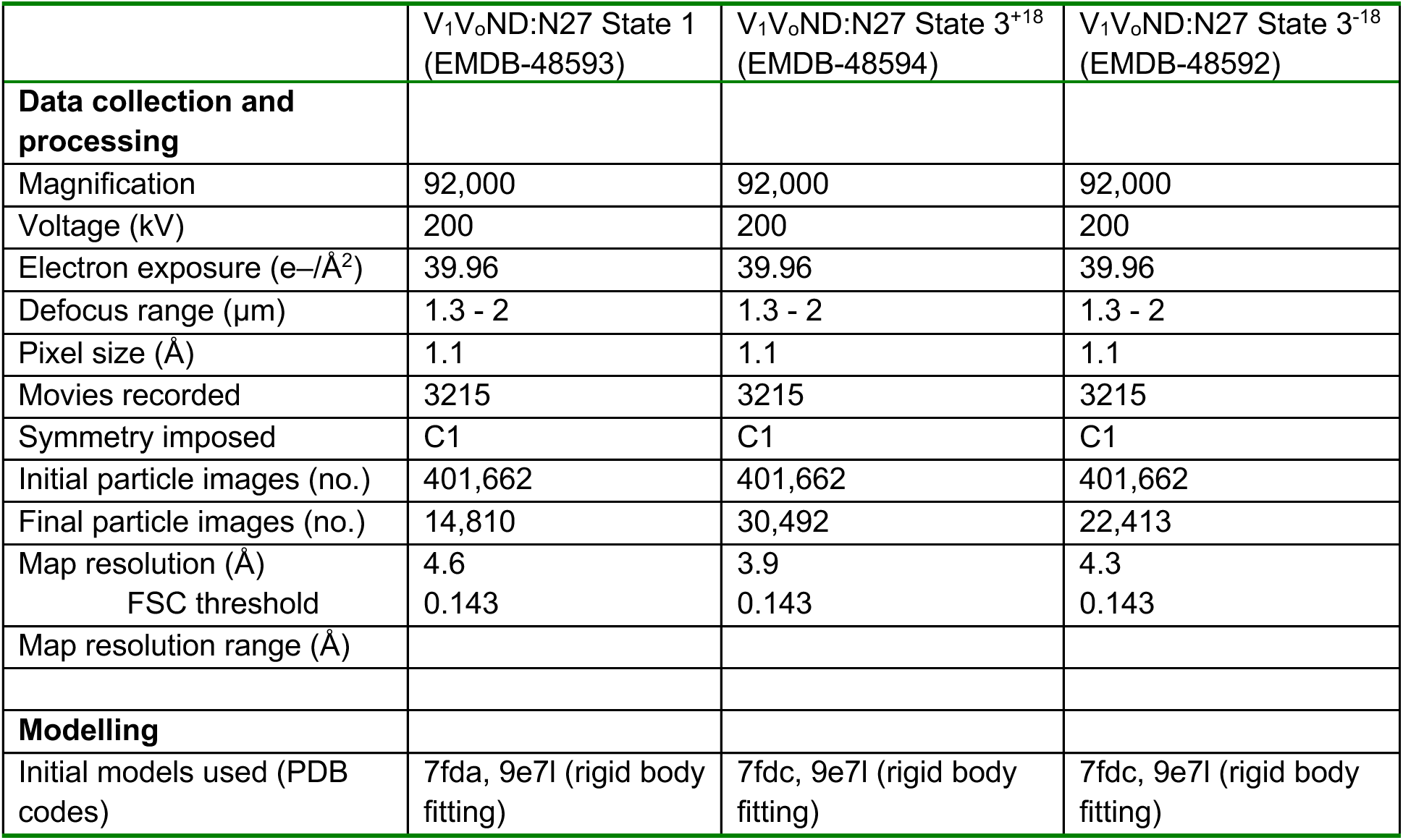
Cryo-EM data collection and modelling of V1Vo:N27 rotary states.

**Appendix Table S3.** Focused refinement of Vo:N27 from V1Vo:N27 rotary states.

|  | V<sub>o</sub>ND:N27 State 1<br>(EMDB-48595) | V<sub>o</sub>ND:N27 State 3<sup>+18</sup><br>(EMDB-48597) | V<sub>o</sub>ND:N27 State 3<sup>-18</sup><br>(EMDB-48596) |
| --- | --- | --- | --- |
| **Data collection and processing** |  |  |  |
| Magnification | 92,000 | 92,000 | 92,000 |
| Voltage (kV) | 200 | 200 | 200 |
| Electron exposure (e<sup>-</sup>/Å<sup>2</sup>) | 39.96 | 39.96 | 39.96 |
| Defocus range (μm) | 1.3 - 2 | 1.3 - 2 | 1.3 - 2 |
| Pixel size (Å) | 1.1 | 1.1 | 1.1 |
| Symmetry imposed | C1 | C1 | C1 |
| Initial particle images (no.) | 401,662 | 401,662 | 401,662 |
| Final particle images (no.) | 14,810 | 30,492 | 22,413 |
| Map resolution (Å) | 4.5 | 3.8 | 4.3 |
| FSC threshold | 0.143 | 0.143 | 0.143 |
| Map resolution range (Å) |  |  |  |
| **Modelling** |  |  |  |
| Initial model used (PDB code) | 7fda, 9e7l (rigid body fitting) | 7fdc, 9e7l (rigid body fitting) | 7fdc, 9e7l (rigid body fitting) |

